# Antigen and Checkpoint Receptor Recalibration of T Cell Receptor Signal Strength

**DOI:** 10.1101/2021.03.02.431957

**Authors:** Thomas A.E. Elliot, Emma K. Jennings, David A.J. Lecky, Natasha Thawait, Adriana Flores-Langarica, David C. Wraith, David Bending

**Affiliations:** Institute of Immunology and Immunotherapy, College of Medical and Dental Sciences, University of Birmingham, Birmingham, B15 2TT, UK; Flow Cytometry Technical Specialist, Infrastructure and Facilities, College of Medical and Dental Sciences, University of Birmingham, Birmingham, B15 2TT, UK

## Abstract

How T cell receptor (TCR) signal strength modulates T cell function and to what extent this is modified by immune checkpoint blockade (ICB) are key questions in immunology. Using Nr4a3-Tocky mice as a digital read-out of NFAT pathway activity, we identify the rapid quantitative and qualitative changes that occur in CD4^+^ T cells in response to a range of TCR signalling strengths. We demonstrate that the time and dose dependent programming of distinct co-inhibitory receptors rapidly re-calibrates T cell activation thresholds. By developing a new *in vivo* model, we analyse the immediate effects of ICB on T cell re-activation. Our findings reveal that anti-PD1 but not anti-Lag3 immunotherapy leads to an increased TCR signal strength. We define a strong TCR signal metric of five genes specifically upregulated by anti-PD1 in T cells (TCR.strong), which can stratify clinical outcomes during anti-PD1 monotherapy in melanoma patients. Our study therefore reveals how analysis of TCR signal strength – and its manipulation – can provide powerful metrics for monitoring outcomes to immunotherapy.

**Key Points:**

- TCR signal strength-dependent programming of CD4^+^ T cells revealed over time in vivo
- Inhibitory receptor expression is dynamic, TCR signal strength dependent, and rapidly re-calibrates T cell activation thresholds
- PD1 but not Lag3 blockade leads to a unique and increased TCR signal strength signature (coined TCR.strong)
- TCR.strong metric stratifies melanoma patient survival in response to Nivolumab (anti-PD1) therapy

## Introduction

How T cells interpret T cell receptor (TCR) signals to promote different functional programmes is a critical aspect of their biology. A key feature of T cell activation is the release of intracellular calcium stores to trigger calcineurin-mediated activation of nuclear factor of activated T cells (NFAT) (Hogan et al., 2003). This process is proposed to occur in a digital and probabilistic fashion (Gallagher et al., 2018; Podtschaske et al., 2007). Similar results are reported for extracellular signal-regulated kinase (ERK) activation (Altan-Bonnet and Germain, 2005; Das et al., 2009). Nonetheless, despite these digital behaviours, it is reported that TCR signal strength can lead to graded expression of molecules such as interferon regulatory factor 4 (IRF4) (Conley et al., 2020), Nr4a1 (Moran et al., 2011) and co-inhibitory receptors (Trefzer et al., 2021). Recent data show that under reduced TCR signal strength levels, nuclear factor kappa B (NF-κB) activation may be analogue (Gallagher et al., 2020). This is intriguing, as NF-κB activation plays critical roles in driving T cell activation, with the activity of mucosa-associated lymphoid tissue lymphoma translocation protein 1 (Malt1) paracaspase is key to full NF-κB activity and IL-2 expression (Rebeaud et al., 2008).

CD8^+^ T cell studies have reported that TCR signal strength does not influence their end stage cytolytic capacity *in vitro* (Richard et al., 2018). However, analysis of thymic CD4^+^ T cell development clearly demonstrates that strong and persistent TCR signals drive Foxp3^+^ Treg development (Bending et al., 2018b; Jennings et al., 2020; Moran et al., 2011), and antigen affinity and dose have distinct effects on peripheral CD4^+^ T cell transcriptional programmes (Keck et al., 2014; Trefzer et al., 2021). Understanding how graded responses to TCR signal strength can modulate T cell function is likely to be critical to understanding mechanisms behind immunotherapies. For example, key T cell transcripts may require differing thresholds of TCR signal strength, meaning some are more amenable to manipulation through immune checkpoint blockade (Shimizu et al., 2020).

Whilst many studies have sought to investigate TCR signalling using *in vitro* systems, the study of how TCR signal strength temporally regulates T cell activation and how immunotherapy may alter these processes are far from clear. In particular, antigen levels can influence the rates of T cell activation, meaning that *in vivo* studies may struggle to dissect differences that occur due to differing T cell activation kinetics (Richard et al., 2018). Furthermore, different T cell genes may require differing durations of TCR signals for expression (Jennings et al., 2020).

To address the challenges of studying T cell activation dynamics, we previously developed the Nr4a3-Timer of cell kinetics and activity (Tocky) model (Bending et al., 2018b). Nr4a3-Tocky mice are NFAT-responsive distal TCR signalling reporter mice (Jennings et al., 2020; Jennings et al., 2021), that exhibit reduced background expression compared to other TCR signalling reporters. Nr4a3-Tocky utilises a fluorescent timer protein (Subach et al., 2009) to monitor the temporal dynamics of TCR signalling and can classify TCR signals according to whether they are *new, persistent,* or *arrested*. Importantly, given that NFAT is necessary and sufficient for expression of Nr4a3 in T cells (Jennings et al., 2020; Martinez et al., 2015), we predicted that Nr4a3 would represent a digital read-out for T cell activation *in vivo*, which would permit the tracking of T cells following antigen encounter over the first 24 hours. Here we employ the Tg4 Nr4a3-Tocky (Jennings et al., 2020) mouse model to track synchronised T cell activation *in vivo*, and identify quantitative and qualitative changes that occur in T cells receiving different strengths of TCR signalling *in vivo*. Importantly this system accounts for differing proportions of T cells that may respond whilst also permitting the analysis of T cells at different synchronised phases following T cell activation. Our findings identify the relationships between TCR signal strength and key T cell transcriptional programmes, including the programming of temporally distinct co-inhibitory receptor modules. These modules rapidly re-calibrate the activation threshold of T cells, allowing direct detection of the immediate effects of immune checkpoint blockade on the re-activation of T cells *in vivo*. Our findings reveal that anti-PD1 but not anti-Lag3 immunotherapy leads to an increased TCR signal strength signature. We refine a TCR signal strength metric down to 5 genes specifically upregulated by anti-PD1 in T cells (called TCR.strong), which stratifies clinical outcomes following anti-PD1 monotherapy in melanoma patients.

## Results

### Antigen dose drives digital Nr4a3 activation at the single cell level but graded responses at population and phenotypic levels

To investigate the regulation of T cell activation, we utilised the Nr4a3-Tocky system, which is an NFAT-dependent distal TCR signal reporter (Jennings et al., 2020) crossed with the Tg4 TCR transgenic line that recognises myelin basic protein (MBP) peptide (Figure 1A). This system has been useful in assessing the response of T cells to modified self-antigens, and it is known that repeated dosing of this system imparts a Type 1 regulatory (Tr1) phenotype (Burton et al., 2014) supported by epigenetic remodelling (Bevington et al., 2020). Therefore, to monitor changes in IL-10 expression our model also incorporated an IL-10-IRES-GFP reporter (Kamanaka et al., 2006). *In vitro* experiments demonstrated that activation with the native lysine at position 4 [4K] MBP peptide induced weak activation of Tg4 T cells (Supplementary Figure 1A&B). Through switching of the fourth peptide residue to alanine [4A] or tyrosine [4Y], the potency of TCR signalling increased significantly (Supplementary Figure 1A&B).

**Figure 1:**
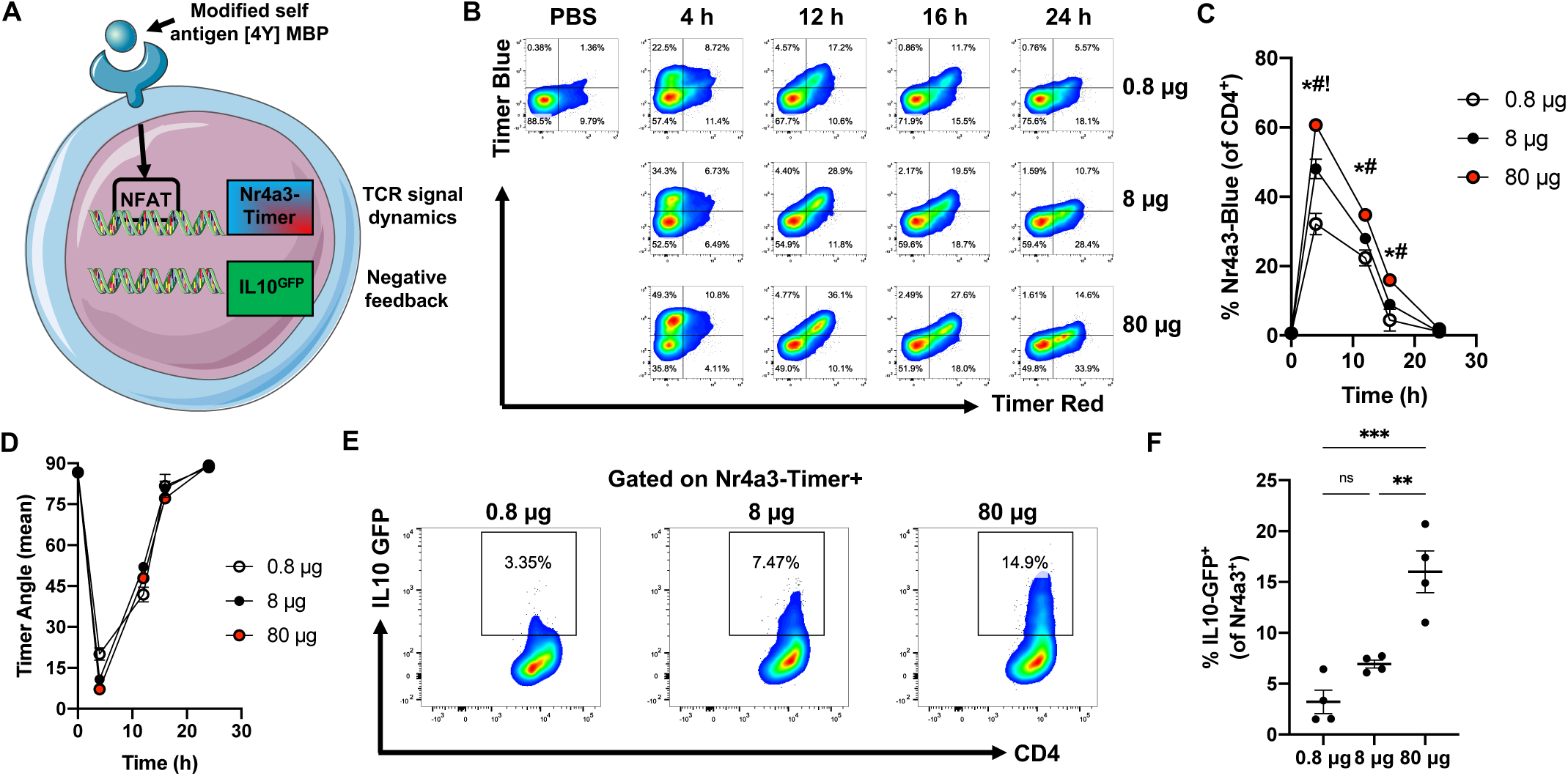
Antigen dose drives digital Nr4a3 activation at the single cell level but graded responses at population and phenotypic levels. (**A**) Schematic detailing the Tg4 Nr4a3-Tocky IL10-GFP system. (**B**) Tg4 Nr4a3-Tocky IL10-GFP mice were immunised s.c. with 0.8, 8 or 80 μg of [4Y] MBP peptide and splenic CD4^+^ T cell responses analysed for Nr4a3-Timer Red vs. Nr4a3-Timer Blue expression in live CD4^+^ Tg4 T cells at the indicated time points. (**C**) Summary data of the % of CD4^+^ Tg4 T cells exhibiting active TCR signalling (% Nr4a3-Blue^+^) or (**D**) mean Nr4a3-Timer Angle at the indicated time points in 0.8 μg (white) 8 μg (black) or 80 ug (red) immunised mice. Bars represent mean±SEM. Statistical analysis by two-way Anova with Tukey’s multiple comparisons test. Significant differences (p<0.05) between 80 μg and 0.8 μg (*), 80 μg and 8 μg (^#^) or 8 μg and 0.8 μg (^!^). (**E**) Tg4 Nr4a3-Tocky IL10-GFP mice were immunised s.c. with 0.8, 8 or 80 μg of [4Y] MBP peptide and splenic CD4^+^ T cell responses analysed for CD4 vs IL10-GFP in Nr4a3-Timer^+^ T cells at 24 h post immunisation. (**F**) Summary data of IL10-GFP expressers (% of CD4^+^) in the three experimental groups. N=4, bars represent mean ±SEM, statistical analysis by one-way Anova with Tukey’s multiple comparisons test. ***=p<0.001, **=p<0.01.

To determine how TCR signal strength affects NFAT/Nr4a3 activation *in vivo*, a hundred-fold range of [4Y] MBP peptide was administered to Tg4 Nr4a3-Tocky IL-10-GFP mice, and Nr4a3 expression monitored over the first 24 h period (Figure 1B-D). As expected at 4 h, splenic T cells responded with an increase in Nr4a3-Blue expression, indicating new TCR signalling in response to recognition of the [4Y] MBP peptide. By 12 h, a population of CD4^+^ T cells had become Nr4a3-Blue^+^Red^+^, indicating the increased time elapsed since initiation of TCR signalling. Strikingly, by 16-24 h responding T cells had migrated towards the *arrested* Nr4a3-Timer locus (Figure 1B) (Bending et al., 2018b). This indicated that the majority of T cells experiencing stimulation in this *in vivo* model arrest Nr4a3 expression within the first 24 h. Analysis of active TCR signalling showed a peak at 4 h, before a fall to near-zero by 24 h (Figure 1C). These results show that the proportion of responding T cells was dependent on TCR signal strength. However, analysis of Nr4a3-Timer Angle (which determines the average position of Nr4a3-Timer^+^ T cells in Blue-Red space, (Bending et al., 2018b)) showed highly similar Timer trajectories independent of the immunising dose (Figure 1D). Therefore, the strength of TCR signalling did not affect the dynamics of Nr4a3 activation; moreover, at the single cell level those T cells that crossed the threshold of activation of the NFAT/Nr4a3 pathway exhibited highly similar dynamics of Nr4a3 expression. However, in the 24 h stimulation period, early expression of IL-10-GFP emerged within Nr4a3-Timer^+^ T cells with a direct correlation to the amount of immunising antigen (Figure 1E-F), reflecting that TCR signal strength can impart rapid phenotypic heterogeneity within activated Nr4a3^+^ T cell populations *in vivo*.

### CD4^+^ T cells rapidly discriminate stimulation strength through transcriptionally distinct and time-dependent activation profiles

The previous data support that the Tg4 Nr4a3-Tocky system can be used to study the temporal dynamics of T cell activation and differentiation during the first 24 h *in vivo*. Based on the link between IL-10 GFP and TCR signal strength, we hypothesised that signal strength imparts both quantitative and qualitative changes, specifically that the TCR signal strength controls the proportion of activated T cells and phenotypically distinct activation profiles. We repeated *in vivo* subcutaneous immunisations of Nr4a3-Timer Tg4 Tiger mice with [4Y] MBP peptide at a 100-fold dose range (0.8 μg and 80 μg) (Figure 2A). In order to control for quantitative differences between the two conditions and identify purely qualitative responses to signal strength, we sorted cells based on their Timer protein maturation profile (4 h Blue^+^Red^-^ *new* signalling, 12 h Blue^+^Red^+^ *persistent* signalling, 24 h Blue^-^Red^+^ *arrested* signalling). This allowed us to isolate T cell populations from different conditions at highly synchronised stages of TCR signalling trajectory, as a comparable period of time had elapsed since the signal threshold for *Nr4a3* expression was met. RNA was extracted from sorted cells and 3’ mRNA libraries were prepared for RNA sequencing. Principal component analysis (PCA) revealed four clusters indicating the control, 4 h, 12 h and 24 h immunised groups (Figure 2B). Within each time cluster, they separated into two distinct groups based on the level of immunising antigen. We decided to focus our analysis on differentially expressed genes (DEGs) between the low and high immunising groups at each time point identified using the DESeq2 algorithm (Love et al., 2014)(Figure 2C, Supplementary Table 1). Our analysis revealed that the highest number of DEGs were present at the 4 h time period, which declined in a time dependent fashion. Analysing the overlap of genes upregulated or downregulated at the different time points suggested that the majority of these genes were unique to the time point of analysis (Figure 2D). Heatmap analysis of the cumulative DEGs across the 3 time points revealed that 24 h samples clustered tightly with the non-activated control population (Supplementary Figure 2A), with 4 h and 12 h clusters separating into discrete branches. To understand biological processes that are influenced by TCR signal strength, we performed Kyoto encyclopaedia of genes and genomes (KEGG) pathway analysis on 4 and 12 h DEGs (Figure 2E-F, Supplementary Table 2). Notable pathways showing significant enrichment at 4 h included, cytokine-cytokine receptor interactions, JAK-STAT signalling, Th17 cell differentiation, Th1 and Th2 cell differentiation and TCR signalling pathways. Interestingly, several of these pathways were still enriched at 12 h (Figure 2F). Analysis of 24 h DEGs reflected sustained changes in cytokine-cytokine receptor interactions and JAK-STAT signalling (Supplementary Figure 2B). Based on these findings, we conceptualised 7 key modules that were undergoing time and dose-dependent transcriptional activation or suppression (Figure 2G). 1) A shared activation module, incorporating the *Nr4a1*-3 receptors, which exhibited rapid induction at 4 h, and was largely off by 12 h. *Cd69* and *Tnf* expression were also tightly linked to this group. 2) A second group of activated genes which trended to have higher and/or longer expression, including TNF superfamily costimulatory receptors *Tnfrsf4* (OX40), *Cd40lg*, the inhibitory receptor *Pdcd1* (PD1), and IL-2 signalling (*Il2* and *Il2ra*). Included here was *Irf4* (previously reported to be graded in response to TCR signalling (Conley et al., 2020)), *Irf8* and *Tbx21* (T-bet). 3) A third module incorporating Th1 associated/ T cell effector functions *(Ifng, Il12rb2, Gzmb)*, which exhibited rapid induction at 4 h in the 80 μg antigen group. Interestingly this module showed delayed activation in the 0.8 μg group; however, *Gzmb* and *Il12rb2* remained high and sustained throughout the 24 h period in 80 μg group. 4) A fourth module specific to strong TCR signalling was upregulated at 4 and 12 h, and largely sustained at 24 h. This included the Th17-associated genes *Rora, Rorc,* and *Il21*. In addition, *Malt1*, an enzyme involved in NF-κB signalling (a pathway that is linked to graded responses to TCR signal strength (Gallagher et al., 2020)) was strongly induced at 4 h and 12 h in 80 μg stimulated group along with the *Mt1* and *Mt2* enzymes involved in zinc bioavailability. 5) The fifth module involved genes undergoing strong and sustained expression that was largely specific to high antigen dose. These included *Ctla4, Icos* and *Maf.* 6) The sixth module identified a delayed and negative regulatory motif which appeared transiently at the 12 h time point in the 80 μg group. This module included *Il10* (echoing the findings in Figure 1E)*, Lag3, Nfil3* and *Tigit* which are all associated with Tr1 cells. Interestingly this module peaked after the termination of TCR signalling (as evidenced by *Nr4a1-3* expression dynamics) in the 80 μg immunised group. A final module exhibited strong but transient downregulation of key parts of the TCR signalling pathway (*Cd3g*, *Cd3e*, *Lck*, *Rasgrp1*) at the 4 h period in the high antigen dose group, indicating negative feedback responses to high antigen dose are stronger in this group *in vivo*; however, by 24 h the expression of these genes had returned to baseline levels. Included in this group was also the Th2-associated transcription factor *Gata3* that was strongly downregulated at 4 h and 12 h, reflecting the KEGG pathway analysis of T cells inducing signatures of Th1 and Th17 programmes (*Rora, Rorc, Tbx21, Ifng, Il12rb2, Il21*). Analysis of DEGs across all time points, revealed that *Il21, Il12rb2, Tbx21, Maf* and *Malt1* were sustained across the whole 24 h period, indicating a motif strongly associated with T cells experiencing a very strong TCR signal *in vivo* (Supplementary Figure 2C). In summary, our analysis identified clear signatures of diverse transcriptional programmes being induced in a time and dose-dependent fashion *in vivo*. These data suggest that, perhaps dissimilarly to reports for CD8^+^ T cells (Richard et al., 2018), early CD4^+^ T cell differentiation fates display greater tunability to antigen levels, and exhibit hallmarks of multiple T helper and T regulatory lineages.

**Figure 2:**
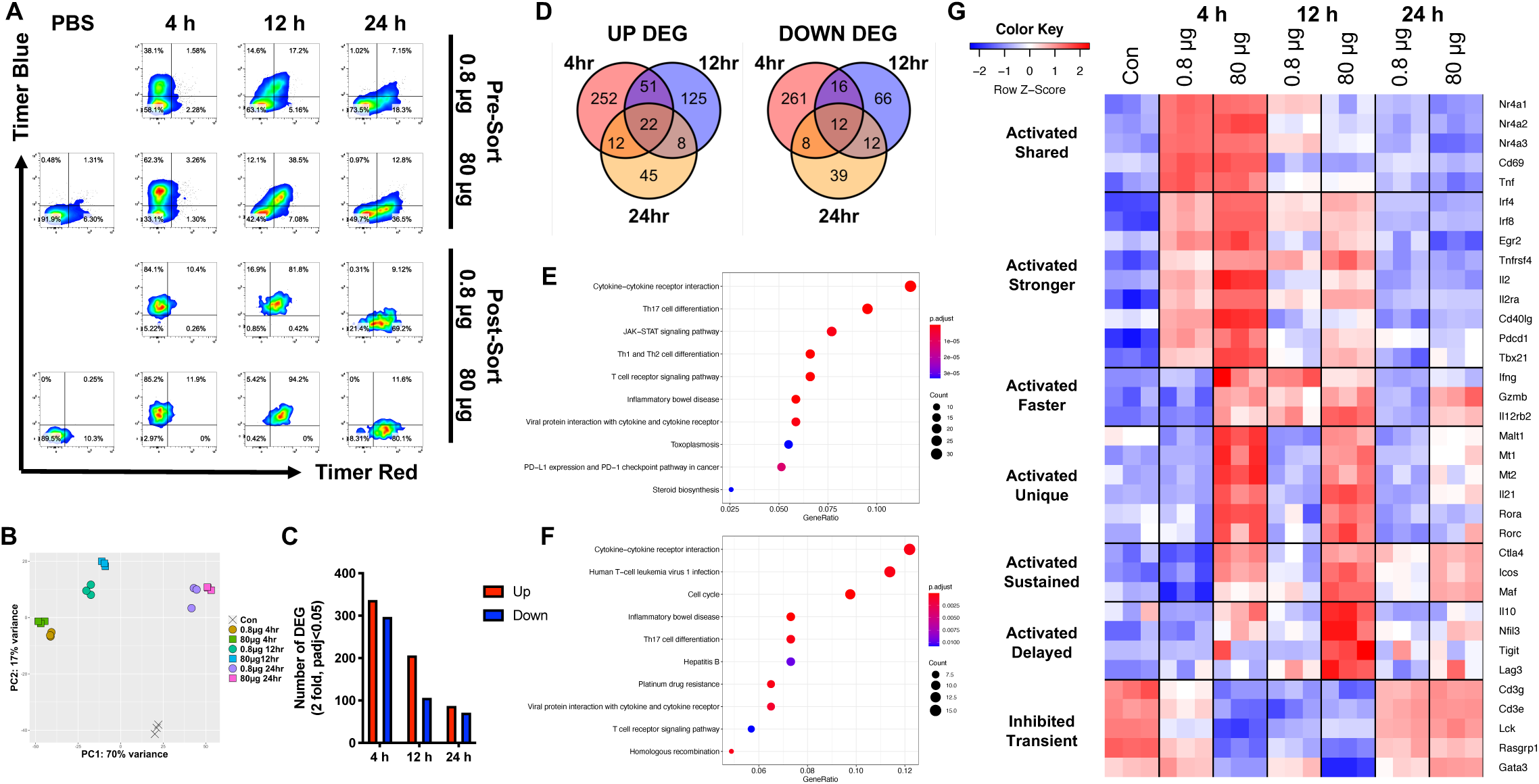
CD4^+^ T cells rapidly discriminate stimulation strength through transcriptionally distinct and time-dependent activation profiles. (**A**) Tg4 Nr4a3-Tocky IL10-GFP mice were immunised s.c. with 0.8 or 80 μg of [4Y] MBP peptide and splenic CD4^+^ T cell responses analysed for Nr4a3-Timer Red vs. Nr4a3-Timer Blue expression in live CD4^+^ Tg4 T cells at the indicated time points (top panel before sorting and bottom panel after cell sorting). (**B**) RNA was extracted from the sorted populations and 3’ mRNA sequencing performed. Principal component analysis of the normalised expression data identifies 7 clusters containing n=3 biological replicates. (**C**) Differentially expressed genes (DEGs) identified using DESeq2 (2-fold change, adjusted p value<0.05) between 80 μg and 0.8 μg stimulated T cells at indicated time points. Up DEG are in red and down DEG in blue. (**D**) Venn diagram analysis of up and down DEG at 4-,12- and 24 h time points. KEGG pathway analysis of DEG between 80 μg and 0.8 μg at 4 h (**E**) or 12 h (**F**) time points. (**G**) Z-score heatmap analysis of log2 transformed and normalised counts of transcripts identifying 7 modules related to the immunising dose and time elapsed since T cell priming.

### *Nr4a3* activation threshold is calibrated by dose dependent negative feedback

A key finding from the time-dependent analysis of transcriptional programmes from T cells activated *in vivo* was the relationship between key negative regulators and TCR signal strength (Figure 3A). PD1 for instance was tightly coupled to T cell activation (Supplementary Figure 3A) and only modestly influenced by TCR signal strength; CTLA-4 (Supplementary Figure 3B) was very much dependent on the immunising dose, and differences in expression were detectable within 4 h. Interestingly the Tr1-associated module exhibited delayed and transient induction in response to strong TCR signalling (Lag3, Tigit and IL10, Supplementary Figure 2D and Figure 1E-F). Given this upregulation of multiple immune checkpoints and/or immunoregulatory cytokines we hypothesised that T cell responsiveness to acute re-stimulation would be dependent on the immunising dose. Moreover, we hypothesised that the T cells that arrested TCR signalling in response to weak TCR signalling would be more sensitive to restimulation than T cells initially activated with a strong TCR signal (Figure 3B). Given that Tg4 Nr4a3-Tocky T cells activated with peptide for 24 h move into the Blue^-^Red^+^ quadrant, due to *arrested* TCR signalling (Figure 1B-D), rechallenge with peptide at this time point would lead to the re-emergence of Nr4a3-Blue expression in this population (Figure 3B), and move up into the Blue^+^Red^+^ quadrant. If the re-challenge duration is restricted to 4 h (Figure 3B), then almost all Nr4a3-Blue^+^Red^+^ T cells will represent T cells that are responding to the first and second stimulations. This is possible due to the fact that the half-life of Nr4a3-Blue protein is 4 h whilst Nr4a3-Red is 120 h (Bending et al., 2018a). T cells that remain in the lower right quadrant (Nr4a3-Red^+^Blue^-^) even after re-stimulation would reflect T cells that fail to re-activate in response to the second dose. To test this hypothesis, we immunised mice with either 0, 8 or 80 μg of [4Y] MBP peptide to induce no, moderate or strong TCR stimulation. 24 h later, we sub-divided these three groups into two groups to receive a further 8 or 80 μg stimulation for 4 h to trigger Nr4a3-Blue expression. Our analysis focused on assessing the proportion of T cells that remained within the *arrested* TCR signalling quadrant (i.e., Nr4a3-Blue^-^Red^+^). Administration of 0 μg followed by 8 μg or 80 μg induced as expected cells predominantly in the *new* Timer locus (Figure 3C, left). Immunising with an initial 8 μg and then rechallenge with 8 μg or 80 μg induced a clear Nr4a3-Blue^+^Red^+^ indicating that the majority of these previously activated T cells responded to the second dose (Figure 3C, middle). In contrast, most T cells immunised with 80 μg and challenged with 8 μg remained in the *arrested* Timer locus (Figure 3C, right). Even when re-stimulating with 80 μg, a significant proportion of *arrested* TCR signalling cells remained. Summary analysis revealed that significantly more T cells failed to re-activate Nr4a3 expression in response to a second restimulation with 8 μg or 80 μg when the T cells had been first immunised with 80 μg (Figure 3D). Importantly this defect was not linked to any differences in the levels of TCR or CD4 (Figure 3E). In summary, our data reveal that T cell activation thresholds are recalibrated by the initial TCR signalling episode *in vivo*.

**Figure 3:**
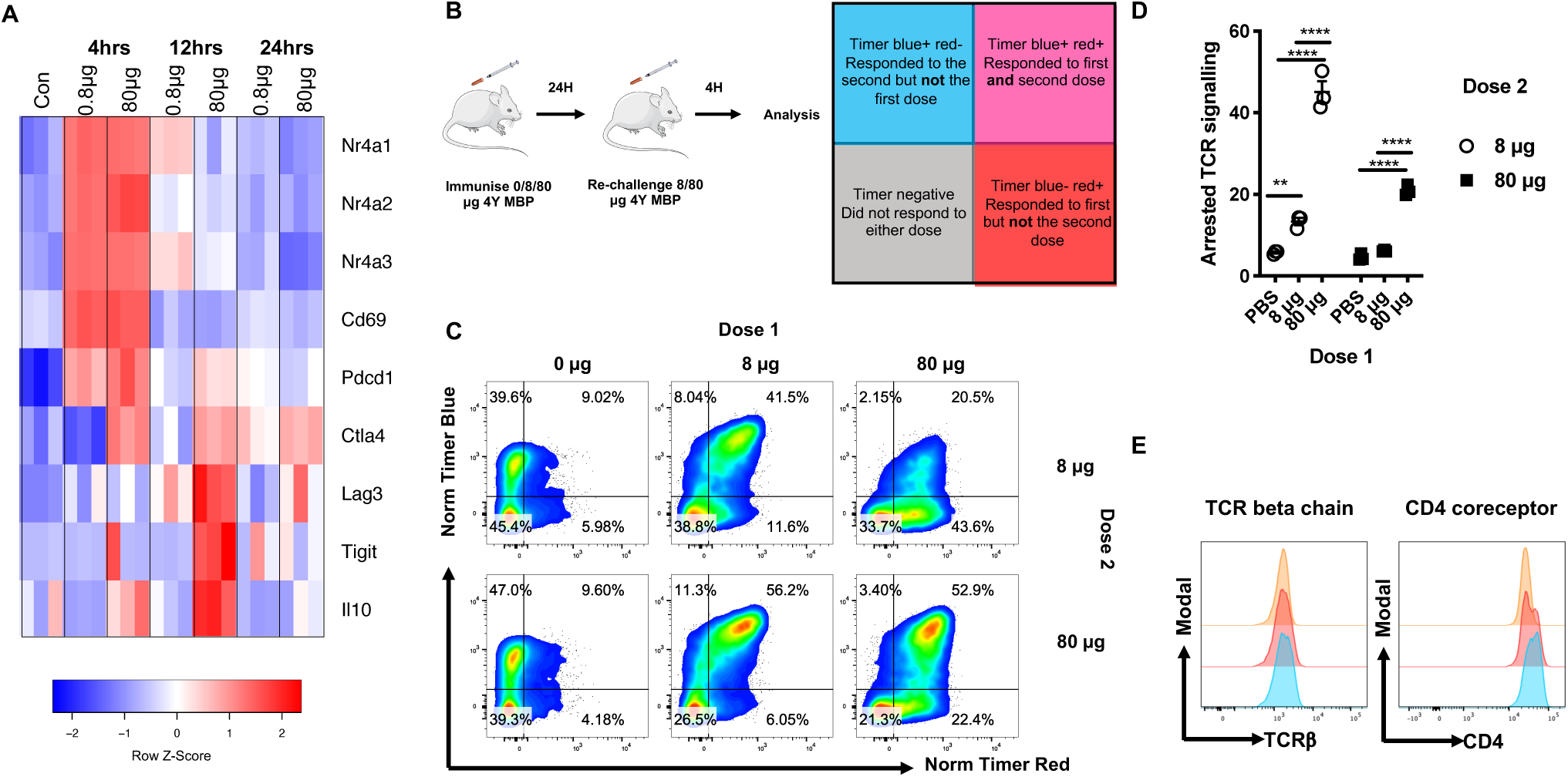
*Nr4a3* activation threshold is calibrated by dose dependent negative feedback. **(A)** Heatmap analysis comparing key inhibitory receptors and their relationships to Nr4a receptor expression from Figure 2G. (**B**) Diagram illustrating experimental setup and interpretation for part (C). (**C**) Tg4 Nr4a3-Tocky IL10-GFP mice were immunised s.c. with 0, 8 or 80 μg of [4Y] MBP. 24 h later mice were randomised to receive either 8 or 80 μg [4Y] MBP rechallenge before splenic CD4^+^ T cells were analysed for Normalised Nr4a3-Timer Blue vs. Normalised Nr4a3-Timer Red analysis 4 h after peptide rechallenge. (**D**) Summary data displaying the frequency of *arrested* (Nr4a3-Blue^-^Red^+^) TCR signalling T cells from (**C**). N=3, bars represent mean ±SEM, statistical analysis by two-way Anova with Sidak’s multiple comparisons test. (**E**) Expression of TCRβ and CD4 on CD4^+^ T cells 28hrs after priming with the indicated dose of [4Y] MBP peptide.

### Co-inhibitory receptors exert distinct quantitative and qualitative control over T cell re-activation

Exposure of self-reactive T cells to a high dose of modified self-antigen led to reduced responsivity to further stimulation and this was correlated with high levels of immune checkpoint receptor expression. We therefore explored the extent to which different checkpoints can modulate the thresholds for re-activation of T cells *in vivo*. We chose PD1, CTLA4/CD28 and Lag3 pathways to compare checkpoints from three modules identified in Figure 3A. We adapted the model from Figure 3B, to include administration of a blocking antibody to the co-inhibitory receptor 30 minutes before the 4 h rechallenge with [4Y] MBP peptide (Figure 4A). In this way we could gain novel insights into the immediate effects of co-inhibitory receptor blockade on the T cell re-activation process *in vivo*. Experiments administering an agonistic CD28 antibody (to bypass potential effects of high CTLA4) did not alter the threshold for activation of T cells in this model (Supplementary Figure 4A&B), we therefore focused our exploration on the potential roles Lag3 and PD1 play in modulating T cell re-activation (Figure 4B). Anti-PD1 blockade induced a significant increase in responders (Figure 4B-D), with anti-Lag3 inducing an intermediate effect on the re-activation of T cells. Interestingly anti-PD1 induced higher levels of Nr4a3-Blue in responding T cells (Figure 4D), than mediated by the isotype pool group or anti-Lag3. These data support that PD1, and to a lesser extent Lag3, quantitatively control the activation thresholds of T cells *in vivo* as reported by NFAT/Nr4a3 activity.

**Figure 4:**
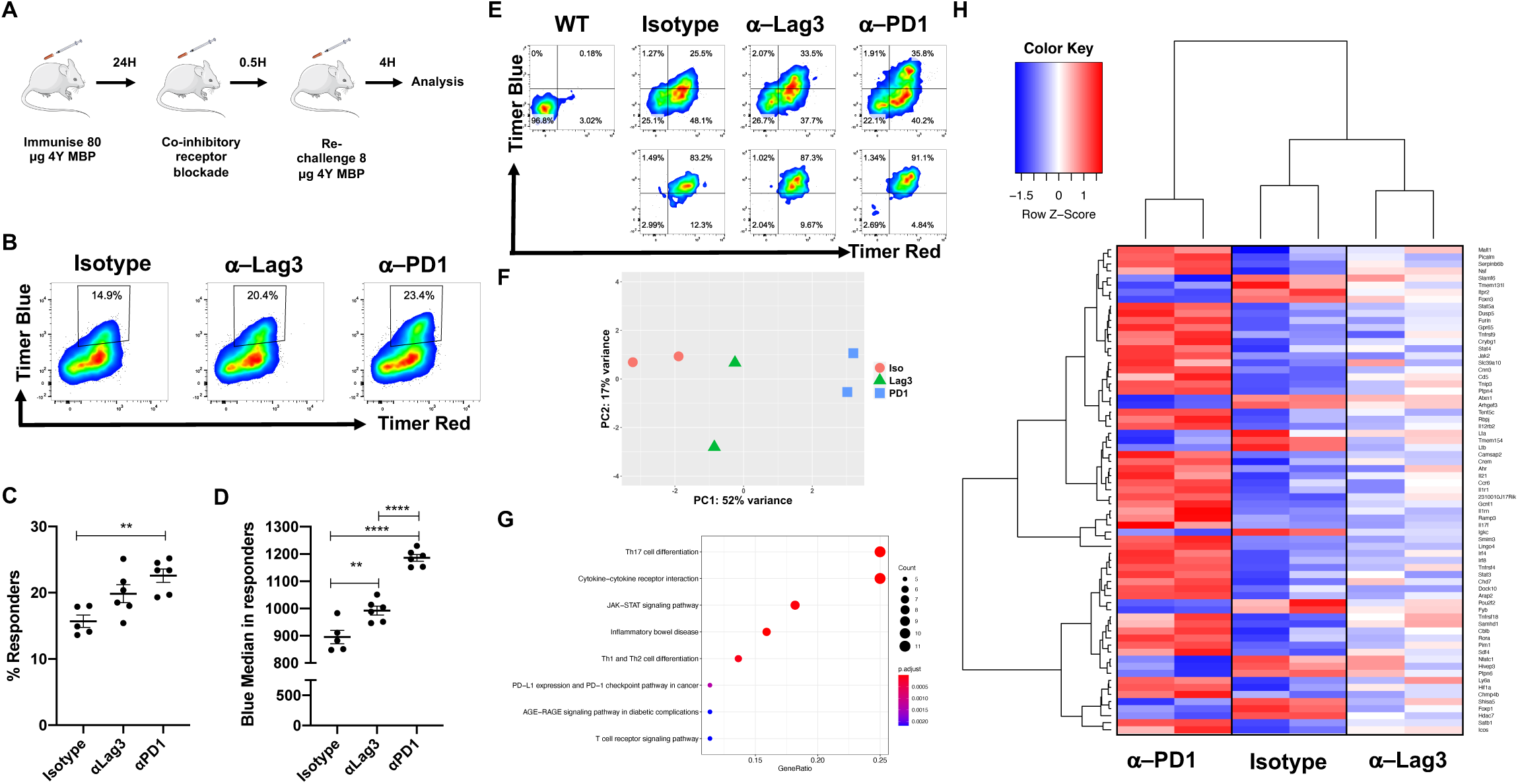
Co-inhibitory receptors exert distinct quantitative and qualitative control over T cell re-activation. (**A**) Diagram detailing experimental design for blockade of co-inhibitory receptors. (**B**) Tg4 Nr4a3-Tocky IL10-GFP mice were immunised s.c. with 80 μg of [4Y] MBP. 24 h later mice were randomised to receive either 0.5 mg isotype pool (1:1 ratio of rat IgG1 and rat IgG2a), anti-Lag3 (clone C9B7W) or anti-PD1 (clone 29F.1A12) 30 minutes prior to re-challenge with 8 μg [4Y] MBP peptide. Splenic CD4^+^ T cells were analysed for Nr4a3-Timer Blue vs. Nr4a3-Timer Red analysis 4 h after peptide rechallenge. Summary data from (**B**) detailing the % Responders (% Nr4a3-Blue^+^Red^+,^ **C**) or median Nr4a3-Blue within Nr4a3-Blue^+^Red^+^ CD4^+^ T cells (**D**) in isotype (n=5), anti-Lag3 (n=6) or anti-PD1 (n=6) treated mice. Bars represent mean ±SEM, dots represent individual mice. Statistical analysis by one-way ANOVA with Tukey’s multiple comparisons test. **=p<0.01, ****=P<0.0001. (**E**) Tg4 Nr4a3-Tocky IL10-GFP mice were immunised s.c. with 80 μg of [4Y] MBP. 24 h later mice were randomised to receive 0.8 mg isotype pool (1:1 ratio of rat IgG1 and rat IgG2a), 0.8 mg anti-Lag3 (clone C9B7W) or 0.8 mg anti-PD1 (clone 29F.1A12) 30 minutes prior to re-challenge with 8 μg [4Y] MBP peptide. Splenic CD4^+^ T cells expression of Nr4a3-Timer Blue vs. Nr4a3-Timer Red 4 h after peptide rechallenge in pre-sorted (top) and sorted (bottom) populations. (**F**) RNA was extracted from the sorted populations and 3’ mRNA sequencing performed. Principal component analysis of the normalised expression data identifies 3 clusters containing n=2 biological replicates. (**G**) KEGG pathway analysis of DEG (adjusted p value<0.05) between isotype and anti-PD1 treated groups. (**H**) Z-score heatmap analysis of log2 transformed and normalised counts displaying the 69 DEG (51 up, 18 down) between isotype and anti-PD1 groups, in isotype, anti-PD1 or anti-Lag3 groups.

As anti-PD1 induced higher levels of Nr4a3-Blue, we hypothesised that anti-PD1 may also induce qualitative changes within T cells re-activating in the presence of its blockade. The precise signalling mechanism of PD1 in an *in vivo* environment is still unclear, with recent *in vitro* studies suggesting that it may preferentially target the CD28 co-stimulatory pathway (Hui et al., 2017). Our data presented here provide evidence that it can also lower the threshold for the NFAT/Nr4a3 pathway on previously activated T cells. In order to compare T cells responding to ICB *in vivo*, we isolated Nr4a3-Blue^+^Red^+^ responder T cells from Isotype, anti-Lag3 or anti-PD1 treated mice (Figure 4E). As before, we isolated Nr4a3-Blue^+^Red^+^ to control for differences in the proportions of responding T cells. This experiment once again re-capitulated the quantitative effects of PD1 and Lag3 blockade on the frequency of responding cells (Supplementary Figure 4C-E). RNA was extracted from these sorted T cells and subjected to 3’ mRNA sequencing. PCA analysis showed that anti-PD1 T cells clustered as a separate group to the isotype and anti-Lag3 groups (Figure 4F). DESeq2 analysis revealed that 69 DEGs existed between the anti-PD1 and Isotype group, demonstrating that 4 h of T cell activation in the presence of anti-PD1 is sufficient to impart qualitative changes within responding T cell populations (Supplementary Table 3). KEGG pathway analysis revealed signatures very similar to those observed in the strong TCR signalling analysis in Figure 2E-F. Cytokine-cytokine receptor, JAK-STAT, Th1/Th2/Th17 differentiation, TCR signalling and PD-L1 expression and PD-1 checkpoint pathway in cancer were significantly enriched terms (Figure 4G). Heatmap analysis showed anti-PD1 clustered distinct from Isotype or anti-Lag3 groups (Figure 4H). Notably we saw an enrichment of costimulatory receptors, including *Tnfrsf4*, *Tnfrsf9*, *Tnfrsf18* and *Icos*. In addition to *Irf4* and *Irf8*, as with 80 μg versus 0.8 μg primary TCR stimulation (Figure 2). Strikingly, *Il21*, *Il12rb2* and *Malt1* were also upregulated in anti-PD1 treated T cells (as had been indicated as sustained markers in Figure 2G). Given the striking similarity between the genes upregulated in the anti-PD1 group compared to controls and those identified in T cells stimulated for 4 h with a high antigen dose we compared the intersect of the DEGs between the two mRNA-seq experiments (Supplementary Figure 5). Our analysis showed that 28/51 of the genes upregulated in T cells re-activated in the presence of anti-PD1 were also genes upregulated in T cells experiencing a strong initial TCR signal for four hours. These data support the notion that anti-PD1 not only increases the probability of a T cell re-activating NFAT/Nr4a3 due to lowering the threshold of activation, but also results in a qualitatively stronger TCR signal, which is not evident in response to anti-Lag3 blockade.

### Strong TCR signalling signatures in tumours of anti-PDL1 treated mice

In order to investigate the relevance of our T cell gene signatures further, we utilised the MC38 colorectal cell line model. This model responds to anti-PDL1 therapy (Efremova et al., 2018), enabling us to explore transcriptional changes occurring in tumours in a polyclonal T cell setting. We injected MC38 tumour cells into the flanks of Nr4a3-Tocky IFNg-YFP mice, to examine the dynamics of tumour development and T cell responses. Tumours increased in weight and volume from day 7 to day 14 (Figure 5A). At days 7, 11 and 14, CD8^+^ TILs were analysed for Nr4a3-Timer, PD1, Lag3 and IFNg expression (Figure 5B-E). The frequency of Nr4a3-Timer^+^ T cells increased over time, as did the development of *arrested* Nr4a3-Timer-Blue^-^Red^+^ T cells (Figure 5C). This was accompanied by a time dependent upregulation in the checkpoints PD1 and Lag3, with approximately 50% of T cells PD1^+^ at day 7, rising to 90% by Day 14 (Figure 5D). A time-dependent increase in IFNg expression was also detected, and increasingly the IFNg-YFP^+^ T cells co-expressed Nr4a3-Timer, indicating that these were recently activated CD8^+^ T cells within the tumour microenvironment (Figure 5E). This confirmed that the MC38 model exhibits hallmarks of chronically stimulated CD8^+^ T cell responses and would serve as a useful model for investigating T cell signatures of response to immune checkpoint blockade. We analysed a previously published 3’ mRNA sequencing dataset performed in mice transplanted with MC38 tumours that were subsequently treated with anti-PDL1 or isotype control (Efremova et al., 2018). We identified 357 genes that were upregulated (>1.5-fold, adjusted p value<0.05) in the dataset, and heatmap analysis showed strong signatures in 2 out of 3 anti-PDL1 treated mice, with an intermediate signature in the third anti-PDL1 treated mouse (Figure 5F). We interrogated the genes identified from our analyses in Supplementary Figure 5 (Upregulated in both 80 μg vs 0.8 μg and anti-PD1 vs Isotype datasets), to visualise expression of these genes in the Efremova *et al*. dataset (Figure 5G, 25 out of 28 detected). These data demonstrated that the majority of signature T cell genes identified in Figure 5 were upregulated (or trended to be upregulated) within the anti-PDL1 treatment group in this model, including *Tnfrsf4*, *Icos*, *Irf8*, *Chmp4b* and *Irf4*. This suggests that our T cell signatures identified through the Tg4 Nr4a3-Tocky model are faithful at discriminating T cell responses in an anti-PDL1-responsive tumour model.

**Figure 5:**
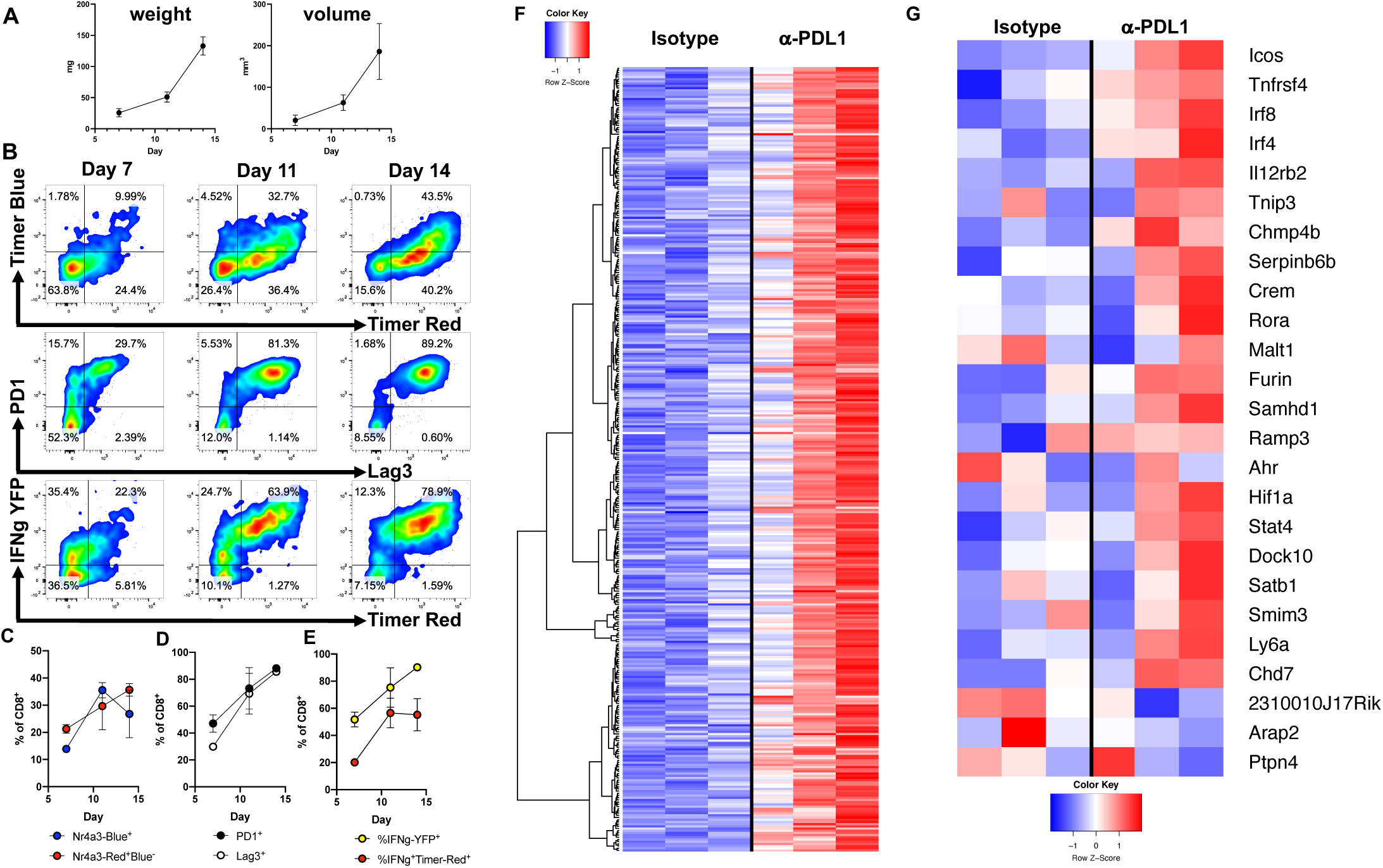
Strong TCR signalling signatures in tumours of anti-PDL1 treated mice. (**A**) 0.25 M MC38 cells were injected s.c. into Nr4a3-Tocky Great Smart-17A mice and tumour weight and volume recorded at days 7, 11 and 14. Bars represent mean ±SEM, n=3. (**B**) CD8^+^ TIL were analysed for Nr4a3-Blue vs Red (top), PD1 vs Lag3 (middle) or Nr4a3-Red vs IFNg-YFP (bottom) expression at the indicated time points. Summary data of % CD8^+^ TIL for (**C**) Nr4a3-Blue^+^ (blue) or Nr4a3-Red^+^Blue^-^ (red), (**D**) PD1^+^ (black) or Lag3^+^ (white), (**E**) IFNg-YFP^+^ (yellow) or %IFNg^+^Nr4a3-Red^+^ (red), n=3 bars represent mean±SEM. (**F**) Z-score heatmap analysis of log2 transformed and normalised counts for genes significantly upregulated (>1.5-fold and adjusted p value<0.05) in C57BL/6 mice injected with 0.5 M MC38 cells then treated with isotype or anti-PDL1 every 3 to four days before whole tumors were excised and 3’ mRNA seq performed (GEO: GSE93018) (Efremova et al., 2018). (**G**) Z-score heatmap analysis of log2 transformed and normalised counts for genes pre-selected from Supplementary Figure 5 and also expressed in GEO: GSE93018 (Efremova et al., 2018).

### Identification of a strong TCR signal metric that stratifies melanoma patient responses to nivolumab therapy

Our findings that anti-PD1 imparts unique signatures of strong TCR signalling led us to hypothesise that genes upregulated in response to strong TCR signalling (Figure 2) or genes upregulated in T cells reactivated in presence of anti-PD1 (Figure 4) could be useful for identifying T cell intrinsic correlates of cancer patients responding to PD1 immunotherapy. We analysed a human gene expression dataset of biopsies from advanced melanoma patients before and after nivolumab therapy. DEGs were examined in those on therapy (**OT**, DEGs Between Pre- and On-therapy Samples, Regardless of Response) or in those with evidence of clinical response (**Res**= DEGs Between Pre- and On-therapy samples, considering genes that change differentially in Responders versus Non-responders) ((Riaz et al., 2017) CA209-038 study). We intersected these genes with DEGs from our Tg4 Nr4a3-Tocky datasets: 4 h T cells stimulated with 80 μg vs 0.8 μg (called **Strong 4hr**) and DEGs in T cells re-activated for 4 h in the presence of anti-PD1 vs isotype (**PD1**). Almost all genes in the on-therapy group were also found within the DEGs in those who exhibited signs of clinical response (Figure 6A, Supplementary Table 4). We identified 5 gene groups of interest intersecting between responders and 4 h/ anti-PD1 (or both). Interestingly, intersection of 4 h and anti-PD1 datasets showed that 2 genes, *ICOS* and *TNIP3,* were associated with both anti-PD1 and strong TCR signalling in mice as well as clinical response to nivolumab, although these genes also changed in patients on therapy regardless of response (group I). This indicates that these may be pharmacodynamic correlates of anti-PD1 therapy. *TNFRSF4* (OX40), *IRF8* and *STAT4* genes were uniquely upregulated only in patients who clinically responded to nivolumab, and also were predicted from our strong 4 h TCR and murine anti-PD1-specific T cell signatures (Figure 6A, group II). This suggests that these transcripts may act as genetic biomarkers of an effective anti-cancer T cell response to therapy. Genes such as *IFNG*, *GZMB* (T cell effector cytokines) and *CTLA4* (checkpoint) were upregulated in the 4 h strong TCR signalling group but also in both the clinical responders and on therapy groups, suggesting that these also show pharmacodynamic responses (group III). Further analysis between clinical response and strong TCR signalling in murine T cells showed that genes associated with immune regulation (*IL2RA*, *IL10*) were significantly upregulated in T cells exhibiting strong TCR signals and only in those patients with objective responses to nivolumab (group IV). Furthermore, we identified *CD5*, *GPR65* and *GCNT1* as a motif that is upregulated on T cells in response to anti-PD1 blockade in mice as well as only in melanoma patients who respond to nivolumab (group V).

**Figure 6:**
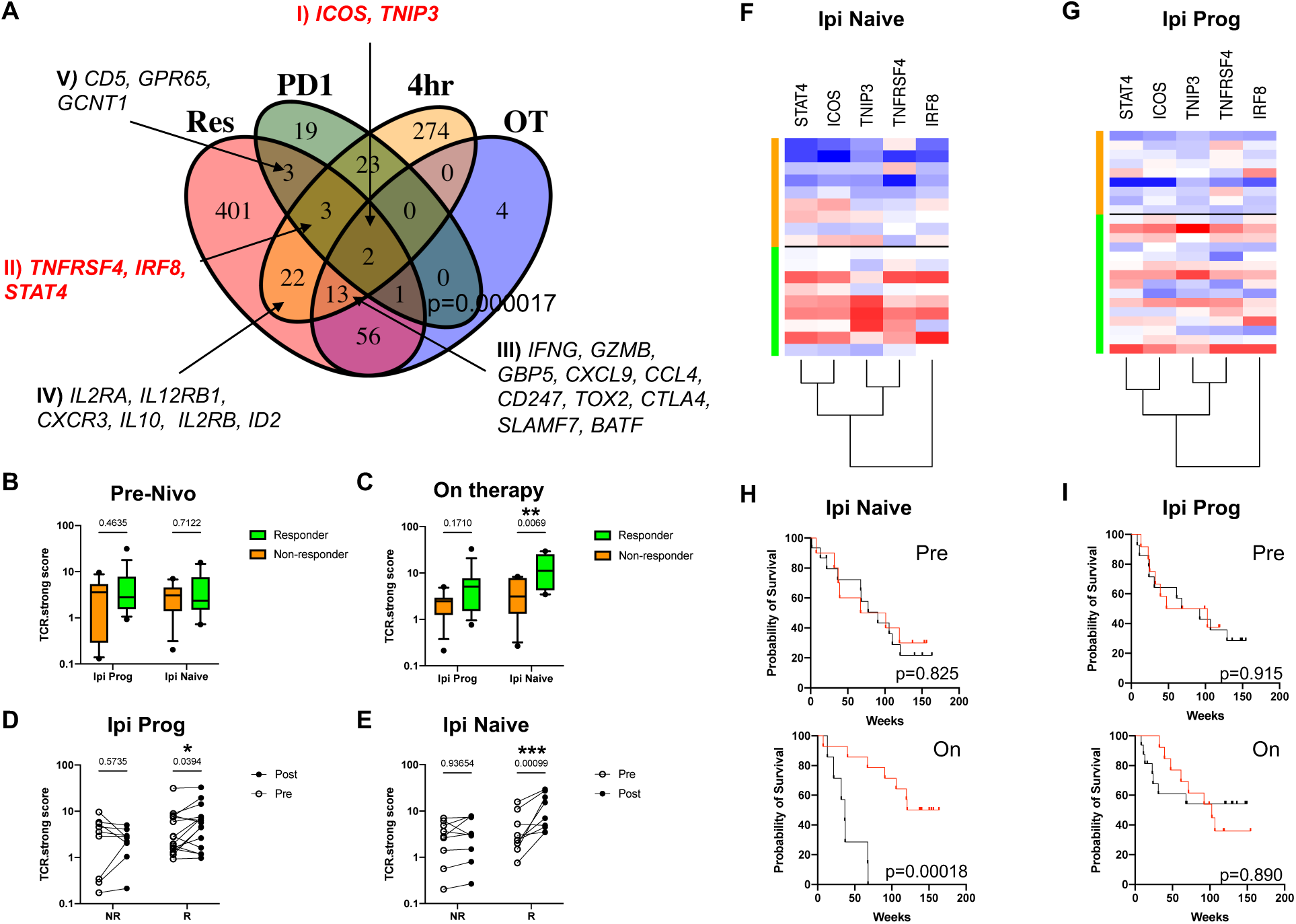
Identification of a strong TCR signal metric that stratifies melanoma patient responses to nivolumab therapy. (**A**) DEG from 4 h time point of 80 μg vs 0.8 μg [4Y] MBP peptide (**4hr**; Figure 2), Isotype versus anti-PD1 (**PD1**; Figure 4) were intersected with DEG from melanoma patients who received nivolumab therapy (Riaz et al., 2017). The DEG in these patients were then classified based on i) the change in expression that occurred on therapy regardless of response compared to pre-therapy samples (**OT**); and ii), DEG that changed compared to pre-therapy samples in those patients showing clinical responses (**Res**). For human datasets a log fold change>0.5 and adjusted p value <0.1 was set. Genes of interest within the sets are annotated. Full lists of genes upregulated in the four datasets are listed in Supplementary Table 4. (**B**) TCR.strong scores in pre-therapy samples in responder (green, ipilimumab (ipi) progressed (Prog) n=15, ipi naïve n=11) and non-responder (orange, ipi prog n=11, ipi naïve n=12). (**C**) TCR.strong scores in on-therapy samples in responder (green, ipi prog n=18, ipi naïve n=10) and non-responder (orange, ipi prog n=11, ipi naïve n=10) patients. (**D**) TCR.strong scores in ipilimumab progressed patients before and after therapy (paired samples indicated by lines), in non-responder (NR, n=9) or responder (R, n=15). (**E**) TCR.strong scores in ipilimumab naive patients before and after therapy in non-responder (NR, n=9) or responder (R, n=9). Bars represent median and whiskers reflect 10 – 90^th^ percentiles in (**B**) and (**C**). Dots represent individual patients and lines pairing of samples in (**D**) and (**E**). Statistical analysis by two-way ANOVA with Sidak’s multiple comparisons test. Z-score heatmap analysis of log2 transformed and normalised counts for TCR.strong metric genes in ipilimumab naïve (**F**) or progressed (**G**) patients. Orange indicates non-responder patients, green indicates responder. Kaplan Meier survival curves for melanoma patients in ipilimumab naïve (**H,** pre-therapy TCR.strong high n=10, TCR.strong low n=15; post-therapy TCR.strong high n=14, TCR.strong low n=7) or ipilimumab progressed patients (**I,** pre-therapy TCR.strong high n=12, TCR.strong low n=14; post-therapy TCR.strong high n=13, TCR.strong low n=16), stratified by high TCR.strong score (>3.5, red) or low TCR.strong score (<3.5, black). Comparison of survival curves by Log Rank test.

Based on these findings, we selected genes in group I (*ICOS*, *TNIP3*) and group II (*TNFRSF4*, *IRF8*, *STAT4*) as the basis for creating a transcriptional signature metric for strong TCR signalling (TCR.strong). We reckoned that by combining indicators of T cell pharmacodynamic responses to anti-PD1 (*ICOS*, *TNIP3*), with genes uniquely changing in patients showing clinical benefit that are also associated with strong TCR signalling and anti-PD1-specific T cell changes (*TNFRSF4, IRF8, STAT4*) we could develop a metric to stratify patient responses to therapy. We utilised an analogous approach to that taken for the Cytolytic score metric, where the geometric mean is taken for the transcripts per million (TPM) from sequencing data *(Roone*y et al., 2015). Using our TCR.strong metric, we interrogated the dataset for nivolumab patients based on their responses and ipilimumab status (Supplementary Table 5). We followed the characterisation of (Riaz et al., 2017), who characterised responding patients (or patient benefit) as those showing complete response, partial response or stable disease. Non-responders (or no-benefit) were classified as those with progressive disease. We divided patients into ipilimumab progressed or naïve, and analysis of the TCR.strong metric in pre-nivolumab biopsies showed no significant differences between responder and non-responders in either ipilimumab progressed or naïve groups (Figure 6B). Analysing biopsies from patients on therapy revealed a strong increase in the metric for the ipilimumab naive cohort in patients with clinical responses (p=0.0069), and a clear but not significant trend in patients who were ipilimumab progressed (p=0.1710, Figure 6C).

To determine the change in the TCR.strong metric before and after therapy, we identified all patients with pre and on therapy biopsies who had known clinical outcomes (Figure 6D-E, Supplementary Table 5). In both the ipilimumab progressed and naïve cohorts, non-responders had no change in their TCR.strong metric, suggesting that the TCR.strong metric is not influenced by anti-PD1 pharmacodynamics. However, ipilimumab progressed patients with clinical responses displayed a small but significant increase (p=0.0394) in their TCR.strong metric (Figure 6D) from pre-therapy levels. For the ipilimumab naïve cohort this increase was more striking and statistically very significant (p=0.00099) in those with evidence of clinical benefit. These differences were clearly illustrated by heatmap analysis of the log2 transformed expression of the 5 genes, showing a stronger and more consistent pattern in the ipilimumab naïve group (Figure 6F) compared to ipilimumab progressed patients (Figure 6G). By using a TCR.strong score threshold of 3.5, we divided patients into TCR.strong high and TCR.strong low groups (Figure 6H-I). Based on this, survival curve analysis showed that high pre-therapy TCR.strong scores had no predictive power in either cohort. In contrast, large survival benefits were found in patients on-therapy with high TCR.strong scores within the cohort of ipilimumab naïve patients (median survival 142 weeks versus 36.6 weeks, p=0.00018). Interestingly, no difference was observed in the ipilimumab progressed groups, suggesting that the mechanism of action of anti-PD1 differs in patients pre-treated with anti-CTLA4. Taken together these data highlight how insights from murine models into transcriptional programmes induced either by strong TCR signalling or PD1 blockade during T cell re-activation can identify TCR signalling based metrics to identify signatures of treatment efficacy in melanoma patients on nivolumab monotherapy.

## Discussion

In this study we sought to understand how antigen levels and immune checkpoints modulate the strength of TCR signalling and the early T cell activation process *in vivo*. As the frequency of a given TCR precursor in a polyclonal setting influences its magnitude of response to an antigen (Moon et al., 2007) we took a reductionist, TCR transgenic approach to focus solely on T cells within the same clonal niche. Through manipulating T cell responses to a modified self-antigen, we identified basic immunological mechanisms that drive the re-calibration of T cell activation thresholds and refine a TCR signal strength metric that can monitor melanoma patient responses to nivolumab.

The use of a highly soluble peptide (Ac-[4Y]-MBP) in the Tg4 TCR transgenic model allowed us to make robust analyses of systemic T cell responses, as our model leads to the rapid and synchronised activation of T cells. This meant we were able to follow the activation trajectories, due to maturation of Nr4a3-Timer protein, of peripheral T cells experiencing different strengths of TCR signalling *in vivo*. Our findings identified several facets of T cell activation that appear to be rapidly programmable due to the strength of TCR signals experienced. We provide evidence that strong TCR signalling leads to the early upregulation of multiple T helper pathways within the same clonal niche, with a bias toward pathways associated with Th1 (*Tbx21*, *Irf4, Il12rb2*, *Stat4*, and repressed *Gata3*), Th17 (*Rorc*, *Rora*, *Il21*), and Tr1 cells (*Maf*, *Nfil3*, *Lag3*, *Il10*). Whilst these findings echo the concept that Th1 cell development depends on the strength of TCR signals compared to Th2 (Constant et al., 1995), multiple studies have identified that TCR signal strength plays a key role in directing CD4^+^ T cell differentiation (Tubo and Jenkins, 2014). Our data, however, suggest that great heterogeneity exists in the early T cell response that does not fit to a simplified model of T helper cell differentiation. These findings echo recent reports of gut CD4^+^ T cell programmes displaying a continuum of phenotypes (Kiner et al., 2021).

Our approach allowed us to only compare T cells that had very recently activated the NFAT/Nr4a3 pathway *in vivo* (Jennings et al., 2020), allowing us to control for the relative frequency of responders whilst also comparing them at similar phases following TCR ligation. This approach also allowed us to interpret the kinetics of the activation of key T cell modules. Interestingly, *Gzmb* and *Ifng*, which are hallmark genes of CD8^+^ T cell responses, were primed in both weak and strong TCR signalling conditions but with differing kinetics. These data suggest that, within T cells that cross the NFAT/Nr4a3 activation threshold, some modules are primed regardless of the TCR signal strength, a finding shown clearly for CD8^+^ T cells and their cytolytic capacity *in vitro* (Richard et al., 2018). However, our findings also reflect that the speed and duration with which these modules are activated differ, with evidence of sustained activation in T cells experiencing a strong TCR signal. In addition, some unique signatures of strong TCR signalling were evident – including the Th17-associated programme (*Rora*, *Rorc*, *Il21*), enzymes involved in zinc bioavailability (*Mt1*, *Mt2*, linked to T cell exhaustion (Singer et al., 2016)), and sustained activation of *Malt1*. Malt1 has essential roles in NF-κB activation in T cells (Rebeaud et al., 2008), and it is tempting to speculate that its function is important in switching NF-κB to full and binary activation that is not achieved by less potent TCR ligands (Gallagher et al., 2020).

Our approach also permitted us to have an in-depth analysis of negative regulators of the T cell activation process, including determining the relationships between key immune checkpoints and the strength and duration of TCR signals. It has been proposed previously that self-peptide/MHC abundance may tune the responses of T cells to antigen (Grossman and Paul, 2015), as shown that those with higher levels of Nur77 and CD5 exhibit hall marks of T cell anergy (such as PD1 and Cbl expression (Zinzow-Kramer et al., 2019)). Our approach reveals how TCR signal strength primes key immune checkpoints with relevance to immunotherapy. Notably CTLA-4 was heavily influenced by the strength of TCR signalling, giving a graded response to antigen dose *in vivo*. In contrast, PD1 showed only a very modest reduction in response to weaker TCR signals, highlighting that PD1 is tightly linked to the activation process. Intriguingly Lag3, Tigit and IL-10, appeared as a delayed and transient module, largely unique to stronger TCR signalling groups. Similar observations have been observed in chronic tolerance models (Burton et al., 2014) and in models of persisting antigen (Trefzer et al., 2021).

Our findings reveal that very strong TCR signalling leads to a rapid re-calibration of T cell activation thresholds at 24 h. We believe this is not due to negative feedback regulation of the TCR signalosome (as CD4 and TCR complexes rapidly recover to control levels by 24 h), but a re-wired T cell activation state akin to adaptive tolerance (Chiodetti et al., 2006). Furthermore, this state is dynamic since it can be overcome by increasing the TCR signal strength through increasing the amount of antigen or through the blockade of negative regulators such as PD1 and Lag3. This tuneable activation threshold allowed us to directly compare the potencies of PD1 and Lag3 in controlling the early re-activation of T cells *in vivo*. Our data clearly show that both exert quantitative control on the frequencies of re-activating T cells; however anti-PD1 showed clear qualitative *in vivo* effects, echoing some recent *in vitro* studies (Shimizu et al., 2020). Other recent data have suggested that Lag3 has a complex mechanism of action but it is likely that it at least in part functions through administering inhibitory signals (Maruhashi et al., 2018). Our data support a weak role in controlling NFAT/Nr4a3 pathway activation, with no evidence that it alters the quality of the resulting TCR signal. This is in contrast to PD1, which imparted features of strong TCR signalling on re-activating T cells. Interestingly the unique signature of T cells re-activated in the presence of anti-PD1 re-capitulated many pathways seen in earlier analyses comparing weak and strong TCR signalling. Once again, a bias towards Th1 and Th17-type pathways was evident, and the strong TCR signalling motif of *Il21*, *Malt1*, *Il12rb2*, *Irf4*, *Irf8*, *Tnfrsf4*, *Tnfrsf18* and *Icos* was readily detectable. In fact, 28/51 genes identified in these analyses overlapped with genes upregulated by strong TCR signals. This finding is intriguing given that anti-PD1 has been proposed to target the anti-CD28 co-stimulatory pathway (Hui et al., 2017). In our study, CD28 agonism had no effect on T cell re-activation, suggesting that anti-PD1 also directly functions to target the TCR driven NFAT/Nr4a3 pathway *in vivo*.

Given the transcriptional features of T cells reactivating in the presence of anti-PD1 we interrogated to what extent strong TCR signatures are evident in human datasets of melanoma patients undergoing nivolumab therapy (Riaz et al., 2017). This dataset is particularly powerful as it permits us to compare genes that change on therapy with those that change in only patients who significantly respond, thereby highlighting potential biomarkers to monitor anti-PD1 responses. It has been clearly documented that IFNγ signatures are a key part of the anti-PD1 response (Grasso et al., 2020; Riaz et al., 2017). In addition, many T cell signatures have been reported to be predictive for anti-PD1 response in a variety of tumour types, including cytolytic (Rooney et al., 2015), T cell IFNγ-related mRNA profiles (Ayers et al., 2017), the chemokine *CXCL9* (Chow et al., 2019; House et al., 2020; Litchfield et al., 2021), *CD8A* (Tumeh et al., 2014) and an antagonistic inflammatory phenotype (Bonavita et al., 2020). A large recent meta-analysis concluded that a compound signature involving tumour mutational burden, *CXCL9*, UV, APOBEC and tobacco signatures can identify pan-cancer responses to ICB (Litchfield et al., 2021). Identification of biomarkers of ICB efficacy before treatment commences would be ideal, as patients could be given treatment based on the likelihood that they will respond. However, given that the majority of patients do not respond to ICB (Borcoman et al., 2019; Sharma et al., 2017), ethically such a test would need to have a very high positive predictive value for widespread clinical application in order to avoid the potential denial of patients of life-extending treatments. Our analysis shows that hallmark signatures of strong TCR signalling or signatures imparted in T cells re-activating in the presence of anti-PD1 can stratify the outcomes of patients on nivolumab therapy. Our TCR.strong metric comprised of 5 immunological genes (*TNFRSF4*, *IRF8*, *STAT4*, *TNIP3*, *ICOS* – the latter previously identified as a potential marker for T cell mediated response to anti-PD1 monotherapy in melanoma, (Xiao et al., 2020)) that were indicated to be genes upregulated rapidly in T cells (<4 h) either experiencing a primary strong TCR signal or in T cells re-activated in the presence of anti-PD1. Importantly the TCR.strong metric was not altered in patients without clinical response, suggesting that these genes are less sensitive to potential pharmacodynamic effects of anti-PD1 therapy. It also demonstrates that this metric cannot predict patient responses before the onset of therapy. However, there remains an urgent need to identify markers to monitor treatment efficacy in patients, in order to inform clinical decision making and enhance the implementation of precision immunotherapy (Havel et al., 2019). In addition, we anticipate that as increasing numbers of ICB combinations become available, identifying signatures for treatment monitoring for efficaciousness will become increasingly as important as identifying predictive biomarkers.

Intriguingly, the TCR.strong metric was most increased in ipilimumab naïve patients and this was reflected in its significant correlation with overall survival in this patient cohort. In fact, it was striking that all patients with low TCR.strong scores died within the ipilimumab naïve cohort, compared to 50% survival in the patients with high TCR.strong scores. These data support the conclusion that strengthened and/or sustained TCR signalling in the tumour environment is a mechanistic feature in patients with clinical response to anti-PD1 therapy, but not in patients previously treated with ipilimumab (anti-CTLA4). Pre-exposure to anti-CTLA4 therapy may reduce the impact of anti-PD1 therapy on the T cell activation signature, given that anti-CTLA4 is likely to encourage an immunologically ‘hot’ environment and enhanced negative feedback (Riaz et al., 2017). Whilst these data are exploratory, the data support investigating the utility of TCR.strong metric in monitoring patients receiving anti-PD1 pathway monotherapy in in future patient cohorts and cancer subtypes.

In summary our study provides new insight into how TCR signal strength and its manipulation control the T cell activation process. Co-inhibitory receptors rapidly re-calibrate the activation threshold of T cells, and we demonstrate how anti-PD1 but not anti-Lag3 immunotherapy leads to a strong TCR signal strength signature that is a correlate for survival of melanoma patients on anti-PD1 monotherapy.

## Materials and Methods

### Mice

Nr4a3-Tocky Tg4 IL10-GFP (Tiger) and Nr4a3-Tocky Great Smart-17A mice (Price et al., 2012) were generated as previously described (Jennings et al., 2020). All animal experiments were approved by the local animal welfare and ethical review body and authorised under the authority of Home Office licences P18A892E0A and PP3965017 (held by D.B.). Animals were housed in specific pathogen-free conditions. Both male and female mice were used, and littermates of the same sex were randomly assigned to experimental groups. Nr4a3-Tocky mice are held under MTA with Dr Masahiro Ono (Imperial College London). Great Smart-17A mice are held under MTA with Prof Richard Locksley (UCSF).

### In vitro cultures

Single cell suspensions of splenocytes were generated as previously described (Jennings et al., 2021). Splenocyte preparations were split in half, with half undergoing naïve CD4^+^ T cells isolation using MoJo magnetic bead negative selection kits (BioLegend), and the other half undergoing CD90^+^ cell depletion (BioLegend) according to the manufacturer’s instructions. Naïve T cells were then mixed at a ratio of 1:1 with CD90-depleted splenocytes and stimulated with 1 μM of acetylated [4K] MBP peptide Ac-ASQKRPSQR, or [4A] Ac-ASQARPSQR or [4Y] Ac-ASQYRPSQR (custom products from GL Biochem Shanghai) in 10% FBS (v/v) RPMI containing 1% penicillin/streptomycin (Life Technologies) at 37°C and 5% CO_2_ for the indicated time points.

### Immunisations

Tg4 Nr4a3-Tocky IL10-GFP (“Tiger” (Kamanaka et al., 2006)) mice were immunised through subcutaneous injection of [4Y] MBP peptide (doses stated in figure legends) in a total volume of 200 μL into the flank. For re-challenge experiments, second doses were administered to the contralateral flank in a volume of 200 μL. Mice were then euthanised at the indicated time points, and spleens removed to analyse systemic T cell responses.

### Antibody treatments

For *in vivo* blockade experiments, *in vivo* grade anti-PD1 (clone 29F.1A12, BioLegend, rat IgG2a), *in vivo* grade anti-Lag3 (clone C9B7W BioLegend, rat IgG1) or hamster anti-CD28 (clone 37.51, kind gift from Prof. Anne Cooke, University of Cambridge) were administered through intraperitoneal injection 30 minutes before peptide rechallenge. For anti-PD1 and anti-Lag3 experiments an isotype pool control group was used consisting of a 1:1 ratio of rat IgG1 (clone MAC221, kind gift from Prof Anne Cooke, University of Cambridge) and rat IgG2a (clone MAC219, kind gift from Prof Anne Cooke, University of Cambridge).

### Flow cytometry and cell sorting

For analysis of splenic lymphocytes, single cell suspensions were prepared as described above. Cells were washed once and stained in 96-well U-bottom plates (Corning). Analysis was performed on a BD LSR Fortessa X-20 instrument. The blue form of the Timer protein was detected in the blue (450/40 nm) channel excited off the 405 nm laser. The red form of the Timer protein was detected in the mCherry (610/20) channel excited off the 561 nm laser. A fixable eFluor 780-flurescent viability dye (eBioscience) was used for all experiments. The following directly conjugated antibodies were used in these experiments: CD4 Alexa Fluor 700 (Clone RM4-4, BioLegend), TCRβ Alexa Fluor 700 (clone H57-597, BioLegend), CD4 BUV737 (Clone GK1.5, BD Biosciences) TCR Vβ8.1, 8.2 PerCP-eFluor 710 (Clone KJ16-133, Thermofisher), TCR Vβ8.1, 8.2 BUV395 (clone, MR5-2, BD Biosciences) CD4 BUV395 (Clone GK1.5, BD Biosciences), CD8 BUV395 (clone 53-6.7, BD Biosciences), TCRβ PerCP-Cy5.5 (clone H57-597, Tonbo Biosciences), PD1 APC (clone 29F.1A12, BioLegend), Tigit PE-Cy7 (clone 1G9, BioLegend), Lag3 APC or PE-Cy7 (clone C9B7W, BioLegend), CTLA-4 PE (clone UC10-4B9, BioLegend), OX40 APC (clone OX-86, BioLegend), GITR PE-Cy7 (clone DTA-1, BioLegend), CD5 APC (clone 53-7.3, BioLegend), ICOS Alexa Fluor 700 (clone C398.4A, BioLegend). For cell sorting, single cell suspensions from biological replicate mice were generated and stained individually with distinct CD4 fluorochromes (e.g. AF700, BUV395, BUV737) to permit multiplexing and parallel cell sorting. Cells were sorted on a FACS Aria cell sorter gating on Nr4a3-Blue^+^Nr4a3-Red^-^ for 4 h time point, Nr4a3-Blue^+^Nr4a3-Red^+^ for 12 h time point and Nr4a3-Blue^-^Nr4a3-Red^+^ for the 24 h time point. For cell sorting in Figure 4, cells were sorted for Nr4a3-Blue^+^Red^+^ T cells. Cells were sorted into 20% FBS RPMI. A small portion of sorted T cells were re-analysed on the flow cytometer to assess purity. Remaining cells were centrifuged for 5 minutes at 500 g before 100µL of extraction buffer added (Arcturus Picopure RNA kit, Thermofisher) and lysates frozen at −80°C.

### MC38 model

MC38 colorectal cell line (kind gift from Prof. David Withers, University of Birmingham) was passaged in 10% FBS (v/v) RPMI containing 1% penicillin/streptomycin (Life Technologies). On day of experiment, MC38 cells were harvested and resuspended in PBS (Sigma) at a concentration of 2.5 million/mL and 0.25 million MC38 cells injected sub cutaneously under the right flank of Nr4a3-Tocky Great Smart-17A mice in a final volume of 100 μL. Tumour size was measured using callipers. Whole tumours from mice were excised, weighed and then dissociated using scissors in 1.2 mL of digestion media containing 1 mg/mL collagenase D (Merck Life Sciences) and 0.1 mg/mL DNase I (Merck Life Sciences) in RPMI. Samples were then incubated for 20-25 minutes at 37°C in a thermoshaker. Digestion mixture was then passed through a 70 μm filter (BD Biosciences) and washed with 30 mL ice cold media (10% FBS RPMI). Suspension was then centrifuged at 1500 rpm for 5 minutes at 4°C. Pellets were then re-suspended in staining media (2% FBS PBS) for labelling with fluorescently conjugated antibodies.

### RNA-seq library preparation and analysis

RNA was extracted from lysates using the Arcturus Picopure RNA kit (Life technologies) according to the manufacturer’s instructions. 15-25 ng of RNA was used for generation of sequencing libraries using the Quantseq 3’ mRNA-seq Library Preparation kit (Lexogen). Briefly, library generation was commenced with oligodT priming containing the Illumina-specific Read 2 linker sequence. After first strand synthesis, RNA was degraded. Second strand synthesis was initiated by random priming and a DNA polymerase. Random primers contained the illumina-specific Read 1 linker sequence. Double stranded DNA was purified from the reaction using magnetic beads and libraries amplified and sequences required for cluster generation and sample indexes were introduced. Libraries were normalised and pooled at a concentration of 4 nM for sequencing. Libraries were sequenced using the NextSeq 500 using a Mid 150v2.5 flow cell. Cluster generation and sequencing was then performed and FASTQ files generated. FASTQ files were then downloaded from the Illumina base space and uploaded to the BlueBee cloud for further analysis (Lexogen). FASTQ files were merged from the 4 lanes to generate final FASTQ files which were loaded into the BlueBee QuantSeq FWD pipeline. FASTQC files were generated and Bbduk v35.92 from the bbmap suite was used for trimming of low-quality tails, poly(A)read-through and adapter contamination. STAR v2.5.2a aligner was used for alignment of reads to the mouse GRCm38 (mm10) genome. HTSeq-count v0.6.0 was used to generate read counts for mRNA species and mapping statistics. Raw read counts in the .txt format were used for further analysis using DESeq2 (Love et al., 2014) in R version 4.0.0. A DESeq dataset was created from a matrix of raw read count data. Data were filtered to remove genes with fewer than 10 reads across all samples. Log2 fold change estimates were generated using the DESeq algorithm and shrinkage using the ashr algorithm (Stephens, 2017) to estimate log2 fold changes. Principal component analysis identified one replicate batch in the effects of checkpoint blockade (Figure 4) to be an outlier and these three samples (which had been sorted and processed as a batch) were not included in further analysis. Differentially expressed genes (DEGs) were selected based on an adjusted p value of <0.05, which was obtained using the Benjamini and Hochberg method for correcting Wald test p values. Normalised read counts were transformed using the regularised log (rlog) transformation. Heatmap analysis was performed on the rlog transformed data using the R package gplots. For KEGG pathway analysis clusterProfiler (Yu et al., 2012), DOSE (Yu et al., 2015), org.Mm.eg.db, and biomaRt (Durinck et al., 2009) packages were used.

### Analysis of human nivolumab and MC38 anti-PDL1 datasets

Genes upregulated in human melanoma patients receiving nivolumab compared to pre-therapy samples were stratified into two groups as in GEO: GSE 91061 (Riaz et al., 2017). On therapy group (**OT**, n=76) consisted of genes upregulated compared to pre-therapy in patients regardless of clinical responses. Responder genes (**Res**, n=501) were those upregulated in patients showing clinical response to treatment compared to pre-therapy. For this analysis a logfold change >0.5 and adjusted p value<0.1 was set. The R package VennDiagram was used for analysis of overlapping genes between the genes in the **OT**, **Res** and genes identified in this study upregulated at 4 h in 80 μg vs 0.8 μg immunised mice (**4hr**, n=337) or genes upregulated 4 h after re-challenge in the presence of anti-PD1 *in vivo* (**PD1**, n=51).

For analysis of gene expression in response to 0.5 mg anti-PDL1 (clone 10F.9G2) therapy compared to IgG2b control in mice inoculated with 0.5 million MC38 cells, raw count expression data was kindly provided by Dr Mirjana Efremova (Efremova et al., 2018) from GEO: GSE93018. Differentially expressed genes (DEG) were identified using DESeq2 as described above. For this analysis genes were considered DEG with a fold change >1.5 and an adjusted p value<0.05. Normalised read counts were transformed using the regularised log (rlog) transformation. Heatmap analysis was performed on the rlog transformed data using the R package gplots.

### Generation and implementation of TCR.strong metric

FPKM for *TNFRSF4*, *ICOS, IRF8*, *TNIP3* and *STAT4* were extracted for nivolumab patients from supplementary files appended to GEO: GSE 91061 (Riaz et al., 2017). FPKM were converted to TPM as described (Pachter, 2011) through dividing each gene level FPKM (offset by 0.01 to deal with 0 counts) by the sum of all FPKM in that sample. This figure was then multiplied by 1^e^6. The geometric mean of the TPM for *TNFRSF4, ICOS, IRF8, TNIP3* and *STAT4* was then calculated, and this value constituted the TCR.strong metric for each patient (Supplementary Table 5). Patient responses were characterised as: complete remission (CR), partial remission (PR), stable disease (SD) or progressive disease (PD) as per (Riaz et al., 2017). Responder groups consisted of CR, PR and SD for analyses. Non-responders were classified as PD patients. Patients with missing disease outcomes or non-evaluated (NE) statuses from supplementary data in (Riaz et al., 2017) were excluded from responder analysis in Figure 6B-E.

### Quantification and statistical analysis

Statistical analysis was performed on Prism 8 or 9 (GraphPad) software. For comparison of more than two means over time, a two-way ANOVA with Tukey’s or Sidak’s multiple comparison’s test was used. For comparison of Kaplan Meier survival curves a Log-rank (Mantel-Cox) test was used. For a comparison of more than two means, a one-way ANOVA with Tukey’s multiple comparisons test was used. Variance is reported as mean ±SEM unless otherwise stated; data points typically represent individual mice. Normalised Nr4a3-Timer Blue, Nr4a3-Timer Red, active TCR signalling and mean Timer angles were generated as previously described using custom algorithms (Bending et al., 2018b). Flow cytometry data were analysed using FlowJo software (BD Biosciences). *p=<0.05, **p=<0.01, ***p=<0.001, ****p=<0.0001.

## Acknowledgements

Work funded by the University of Birmingham (D.B.), the Wellcome Trust (214018/Z/18/Z, to D.B., T.A.E.E, and E.K.J,), and the MRC (MR/V009052/1, to D.B.). D.A.L. is funded by a Wellcome Trust 4-year Basic Science PhD program. E.K.J. is supported by a studentship from the MRC Discovery Medicine North (DiMeN) Doctoral Training Partnership (MR/N103840/1). D.C.W and A.F-L. are funded by the University of Birmingham (UoB). Diagrams in Figures 1A, 3B and 4A were adapted from the Servier Medical Art templates, which are licensed under a CC BY 3.0 unported licence, https://smart.servier.com. We thank Dr Leila Khoja, Dr Neil Steven and Dr Lalit Pallan (Medical Oncology, UoB) for helpful discussion around the potential clinical utility of the TCR.strong metric. We also thank Dr Sarah Dimeloe, Dr Kendle Maslowski, Dr Rebecca Drummond, Dr Wei-Yu Lu and Dr Alastair Copland (all UoB) for their continued support and feedback on data in the manuscript.

## Declaration of interests

The authors declare no competing interests.

## Author contributions

Conceptualisation, funding acquisition, supervision, formal analysis, methodology, data curation, project administration and writing of original draft (D.B.). T.A.E.E, E.K.J., N.T., D.A.J.L and D.B. performed and analysed experiments. D.B. performed and analysed RNA-seq experiments and performed all bioinformatic analyses, including conceptualisation and implementation of the TCR.strong metric. D.C.W. provided resources and advice on methodology for Tg4 model immunisation. A.F.L performed Cell Sorting experiments. All authors were involved in reviewing and editing the original draft manuscript.

## Supplementary Figure and Table Legends

**Supplementary Figure 1: Modified MBP peptide variants induce potent T cell activation *in vitro* (related to Figure 1)**

Naïve CD4+ T cells from Tg4 Nr4a3-Tocky IL10-GFP mice were incubated with CD90-depleted splenocytes in the presence of 1 μM of native [4K] MBP peptide, or [4A] or [4Y] variants for the times indicated before analysis of Nr4a3-Blue vs Nr4a3-Red expression in CD4^+^ Tg4 T cells. (B) Summary data showing the % Nr4a3-Blue^+^ in CD4^+^ T cells in the 3 peptide groups. MBP [4K] = white circles, MBP [4A] = black squares and MBP [4Y] = red circles. Bars represent mean±SEM, n=3. Statistical analysis by two-way ANOVA with Tukey’s multiple comparisons test. Significant differences between [4Y] and [4K] = *, [4A] and [4K] = #, or [4Y] and [4A] = !.

**Supplementary Figure 2: Analysis of DEGs in T cells receiving strong or weak TCR signalling *in vivo* (related to Figure 2).**

**(A)** Z-score heatmap analysis of log2 transformed and normalised counts for all unique DEG identified between 0.8 and 80 μg groups at 4, 12 or 24 h. (**B**) KEGG pathway analysis of DEG at 24 h between 0.8 μg and 80 μg [4Y] MBP immunised mice. (**C**) Z-score heatmap analysis of log2 transformed and normalised counts for genes that show differential expression across all 3 time points (4, 12 and 24 h) analysed between 0.8 μg and 80 μg immunised groups.

**Supplementary Figure 3: Protein expression patterns of PD1, CTLA4, Lag3 and TIGIT in response to different grades of TCR signal strength (related to Figure 3)**

**(A)** Tg4 Nr4a3-Tocky IL10-GFP mice were immunised s.c. with 0.8 μg, 8 μg or 80 μg of [4Y] MBP peptide and splenic CD4^+^ Nr4a3-Timer^+^ T cells (or gated on CD4^+^ PD1^+^ in (**B**)) analysed for PD1 (**A**), CTLA4 (**B**), Tigit (**C**) or Lag3 (**D**) expression. Dots represent individual mice and bars represent mean ±SEM. Statistical analysis by one-way ANOVA with Tukey’s multiple comparisons test.

**Supplementary Figure 4: Effects of CD28, Lag3 and PD1 pathways on T cell re-activation in vivo (related to Figure 4)**

**(A)** Tg4 Nr4a3-Tocky IL10-GFP mice were immunised s.c. with 80 μg of [4Y] MBP. 24 h later mice were randomised to receive either PBS or agonistic anti-CD28 30 minutes prior to re-challenge with 8 μg [4Y] MBP peptide. Splenic CD4^+^ T cells were analysed for Nr4a3-Timer Blue vs. Nr4a3-Timer Red analysis 4 h after peptide rechallenge. (**B**) Summary data of (**A**), control n=4, anti-CD28 n=3. (**C&D**) Tg4 Nr4a3-Tocky IL10GFP mice were immunised s.c. with 80 μg of [4Y] MBP. 24 h later mice were randomised to receive either isotype, anti-Lag3 or anti-PD1 30 minutes prior to re-challenge with 8 μg [4Y] MBP peptide. The frequency of responder (Nr4a3-Blue^+^Red^+^) T cells (**C**) or Nr4a3-Blue Median expression in responder (Nr4a3-Blue^+^Red^+^) T cells (**D**) 4 h after peptide rechallenge are shown. N=3, dots represent individual mice, bars represent mean ±SEM. Statistical analysis by one-way ANOVA with Tukey’s multiple comparisons test. *=p<0.05, **=p<0.01, ***=p<0.001.

**Supplementary Figure 5: Majority of anti-PD1 specific T cell genes are upregulated in T cells receiving strong TCR signal (related to Figures 2 and 4 and 5)**

Genes upregulated in Tg4 CD4^+^ T cells 4 h after receiving 80 μg vs 0.8 μg [4Y] MBP (Figure 2) were intersected with genes selectively upregulated at 4hrs of T cells re-activated in the presence of anti-PD1 *in vivo* (Figure 4). 28 out of 51 genes were overlapping and utilised to interrogate MC38 tumour responses in Figure 5.

## Supplementary Tables

**Supplementary Table 1:** Differentially expressed genes between 80 μg vs 0.8 μg at 4, 12 and 24 h time points (related to Figure 2)

**Supplementary Table 2:** KEGG pathway analysis of DEGs between 80 μg and 0.8 μg groups at 4, 12 and 24 h post immunisation (related to Figure 2)

**Supplementary Table 3:** DEGs between anti-PD1 vs isotype treated responder T cells (related to Figure 4)

**Supplementary Table 4:** Genes upregulated in melanoma patients receiving Nivolumab split according to those upregulated on therapy regardless of clinical response (**OT**), vs those clinically responding (**Re**s), or upregulated in 4 h strong TCR (**4hr,** Figure 2C) or in T cells re-activating in response to anti-PD1 (**PD1,** Figure 4H) (related to Figure 6).

**Supplementary Table 5:** FPKM and TPM for TCR.strong genes, TCR.strong scores, and clinical information for patients analysed (related to Figure 6)

## Notes

### Competing Interest Statement

The authors have declared no competing interest.

## References

Altan-Bonnet, G., and Germain, R.N. (2005). Modeling T cell antigen discrimination based on feedback control of digital ERK responses. PLoS Biol 3, e356.

Ayers, M., Lunceford, J., Nebozhyn, M., Murphy, E., Loboda, A., Kaufman, D.R., Albright, A., Cheng, J.D., Kang, S.P., Shankaran, V., et al. (2017). IFN-gamma-related mRNA profile predicts clinical response to PD-1 blockade. J Clin Invest 127, 2930–2940.

Bending, D., Paduraru, A., Ducker, C.B., Prieto Martin, P., Crompton, T., and Ono, M. (2018a). A temporally dynamic Foxp3 autoregulatory transcriptional circuit controls the effector Treg programme. EMBO J 37.

Bending, D., Prieto Martin, P., Paduraru, A., Ducker, C., Marzaganov, E., Laviron, M., Kitano, S., Miyachi, H., Crompton, T., and Ono, M. (2018b). A timer for analyzing temporally dynamic changes in transcription during differentiation in vivo. J Cell Biol 217, 2931–2950.

Bevington, S.L., Ng, S.T.H., Britton, G.J., Keane, P., Wraith, D.C., and Cockerill, P.N. (2020). Chromatin Priming Renders T Cell Tolerance-Associated Genes Sensitive to Activation below the Signaling Threshold for Immune Response Genes. Cell Rep 31, 107748.

Bonavita, E., Bromley, C.P., Jonsson, G., Pelly, V.S., Sahoo, S., Walwyn-Brown, K., Mensurado, S., Moeini, A., Flanagan, E., Bell, C.R., et al. (2020). Antagonistic Inflammatory Phenotypes Dictate Tumor Fate and Response to Immune Checkpoint Blockade. Immunity 53, 1215–1229 e1218.

Borcoman, E., Kanjanapan, Y., Champiat, S., Kato, S., Servois, V., Kurzrock, R., Goel, S., Bedard, P., and Le Tourneau, C. (2019). Novel patterns of response under immunotherapy. Ann Oncol 30, 385–396.

Burton, B.R., Britton, G.J., Fang, H., Verhagen, J., Smithers, B., Sabatos-Peyton, C.A., Carney, L.J., Gough, J., Strobel, S., and Wraith, D.C. (2014). Sequential transcriptional changes dictate safe and effective antigen-specific immunotherapy. Nat Commun 5, 4741.

Chiodetti, L., Choi, S., Barber, D.L., and Schwartz, R.H. (2006). Adaptive tolerance and clonal anergy are distinct biochemical states. J Immunol 176, 2279–2291.

Chow, M.T., Ozga, A.J., Servis, R.L., Frederick, D.T., Lo, J.A., Fisher, D.E., Freeman, G.J., Boland, G.M., and Luster, A.D. (2019). Intratumoral Activity of the CXCR3 Chemokine System Is Required for the Efficacy of Anti-PD-1 Therapy. Immunity 50, 1498–1512 e1495.

Conley, J.M., Gallagher, M.P., Rao, A., and Berg, L.J. (2020). Activation of the Tec Kinase ITK Controls Graded IRF4 Expression in Response to Variations in TCR Signal Strength. J Immunol 205, 335–345.

Constant, S., Pfeiffer, C., Woodard, A., Pasqualini, T., and Bottomly, K. (1995). Extent of T cell receptor ligation can determine the functional differentiation of naive CD4+ T cells. J Exp Med 182, 1591–1596.

Das, J., Ho, M., Zikherman, J., Govern, C., Yang, M., Weiss, A., Chakraborty, A.K., and Roose, J.P. (2009). Digital signaling and hysteresis characterize ras activation in lymphoid cells. Cell 136, 337–351.

Durinck, S., Spellman, P.T., Birney, E., and Huber, W. (2009). Mapping identifiers for the integration of genomic datasets with the R/Bioconductor package biomaRt. Nat Protoc 4, 1184–1191.

Efremova, M., Rieder, D., Klepsch, V., Charoentong, P., Finotello, F., Hackl, H., Hermann-Kleiter, N., Lower, M., Baier, G., Krogsdam, A., and Trajanoski, Z. (2018). Targeting immune checkpoints potentiates immunoediting and changes the dynamics of tumor evolution. Nat Commun 9, 32.

Gallagher, M.P., Conley, J.M., and Berg, L.J. (2018). Peptide Antigen Concentration Modulates Digital NFAT1 Activation in Primary Mouse Naive CD8(+) T Cells as Measured by Flow Cytometry of Isolated Cell Nuclei. Immunohorizons 2, 208–215.

Gallagher, M.P., Conley, J.M., Vangala, P., Reboldi, A., Garber, M., and Berg, L.J. (2020). The Tec kinase ITK differentially optimizes NFAT, NF-κB, and MAPK signaling during early T cell activation to regulate graded gene induction. bioRxiv, 2020.2011.2012.380725.

Grasso, C.S., Tsoi, J., Onyshchenko, M., Abril-Rodriguez, G., Ross-Macdonald, P., Wind-Rotolo, M., Champhekar, A., Medina, E., Torrejon, D.Y., Shin, D.S., et al. (2020). Conserved Interferon-gamma Signaling Drives Clinical Response to Immune Checkpoint Blockade Therapy in Melanoma. Cancer Cell 38, 500–515 e503.

Grossman, Z., and Paul, W.E. (2015). Dynamic tuning of lymphocytes: physiological basis, mechanisms, and function. Annu Rev Immunol 33, 677–713.

Havel, J.J., Chowell, D., and Chan, T.A. (2019). The evolving landscape of biomarkers for checkpoint inhibitor immunotherapy. Nat Rev Cancer 19, 133–150.

Hogan, P.G., Chen, L., Nardone, J., and Rao, A. (2003). Transcriptional regulation by calcium, calcineurin, and NFAT. Genes Dev 17, 2205–2232.

House, I.G., Savas, P., Lai, J., Chen, A.X.Y., Oliver, A.J., Teo, Z.L., Todd, K.L., Henderson, M.A., Giuffrida, L., Petley, E.V., et al. (2020). Macrophage-Derived CXCL9 and CXCL10 Are Required for Antitumor Immune Responses Following Immune Checkpoint Blockade. Clin Cancer Res 26, 487–504.

Hui, E., Cheung, J., Zhu, J., Su, X., Taylor, M.J., Wallweber, H.A., Sasmal, D.K., Huang, J., Kim, J.M., Mellman, I., and Vale, R.D. (2017). T cell costimulatory receptor CD28 is a primary target for PD-1-mediated inhibition. Science 355, 1428–1433.

Jennings, E., Elliot, T.A.E., Thawait, N., Kanabar, S., Yam-Puc, J.C., Ono, M., Toellner, K.M., Wraith, D.C., Anderson, G., and Bending, D. (2020). Nr4a1 and Nr4a3 Reporter Mice Are Differentially Sensitive to T Cell Receptor Signal Strength and Duration. Cell Rep 33, 108328.

Jennings, E.K., Lecky, D.A.J., Ono, M., and Bending, D. (2021). Application of dual Nr4a1-GFP Nr4a3-Tocky reporter mice to study T cell receptor signaling by flow cytometry. STAR Protoc 2, 100284.

Kamanaka, M., Kim, S.T., Wan, Y.Y., Sutterwala, F.S., Lara-Tejero, M., Galan, J.E., Harhaj, E., and Flavell, R.A. (2006). Expression of interleukin-10 in intestinal lymphocytes detected by an interleukin-10 reporter knockin tiger mouse. Immunity 25, 941–952.

Keck, S., Schmaler, M., Ganter, S., Wyss, L., Oberle, S., Huseby, E.S., Zehn, D., and King, C.G. (2014). Antigen affinity and antigen dose exert distinct influences on CD4 T-cell differentiation. Proc Natl Acad Sci U S A 111, 14852–14857.

Kiner, E., Willie, E., Vijaykumar, B., Chowdhary, K., Schmutz, H., Chandler, J., Schnell, A., Thakore, P.I., LeGros, G., Mostafavi, S., et al. (2021). Gut CD4(+) T cell phenotypes are a continuum molded by microbes, not by TH archetypes. Nat Immunol 22, 216–228.

Litchfield, K., Reading, J.L., Puttick, C., Thakkar, K., Abbosh, C., Bentham, R., Watkins, T.B.K., Rosenthal, R., Biswas, D., Rowan, A., et al. (2021). Meta-analysis of tumor- and T cell-intrinsic mechanisms of sensitization to checkpoint inhibition. Cell 184, 596–614 e514.

Love, M.I., Huber, W., and Anders, S. (2014). Moderated estimation of fold change and dispersion for RNA-seq data with DESeq2. Genome Biol 15, 550.

Martinez, G.J., Pereira, R.M., Aijo, T., Kim, E.Y., Marangoni, F., Pipkin, M.E., Togher, S., Heissmeyer, V., Zhang, Y.C., Crotty, S., et al. (2015). The transcription factor NFAT promotes exhaustion of activated CD8(+) T cells. Immunity 42, 265–278.

Maruhashi, T., Okazaki, I.M., Sugiura, D., Takahashi, S., Maeda, T.K., Shimizu, K., and Okazaki, T. (2018). LAG-3 inhibits the activation of CD4(+) T cells that recognize stable pMHCII through its conformation-dependent recognition of pMHCII. Nat Immunol 19, 1415–1426.

Moon, J.J., Chu, H.H., Pepper, M., McSorley, S.J., Jameson, S.C., Kedl, R.M., and Jenkins, M.K. (2007). Naive CD4(+) T cell frequency varies for different epitopes and predicts repertoire diversity and response magnitude. Immunity 27, 203–213.

Moran, A.E., Holzapfel, K.L., Xing, Y., Cunningham, N.R., Maltzman, J.S., Punt, J., and Hogquist, K.A. (2011). T cell receptor signal strength in Treg and iNKT cell development demonstrated by a novel fluorescent reporter mouse. J Exp Med 208, 1279–1289.

Pachter, L. (2011). Models for transcript quantification from RNA-Seq. arXiv.

Podtschaske, M., Benary, U., Zwinger, S., Hofer, T., Radbruch, A., and Baumgrass, R. (2007). Digital NFATc2 activation per cell transforms graded T cell receptor activation into an all-or-none IL-2 expression. PLoS One 2, e935.

Price, A.E., Reinhardt, R.L., Liang, H.E., and Locksley, R.M. (2012). Marking and quantifying IL-17A-producing cells in vivo. PLoS One 7, e39750.

Rebeaud, F., Hailfinger, S., Posevitz-Fejfar, A., Tapernoux, M., Moser, R., Rueda, D., Gaide, O., Guzzardi, M., Iancu, E.M., Rufer, N., et al. (2008). The proteolytic activity of the paracaspase MALT1 is key in T cell activation. Nat Immunol 9, 272–281.

Riaz, N., Havel, J.J., Makarov, V., Desrichard, A., Urba, W.J., Sims, J.S., Hodi, F.S., Martin-Algarra, S., Mandal, R., Sharfman, W.H., et al. (2017). Tumor and Microenvironment Evolution during Immunotherapy with Nivolumab. Cell 171, 934–949 e916.

Richard, A.C., Lun, A.T.L., Lau, W.W.Y., Gottgens, B., Marioni, J.C., and Griffiths, G.M. (2018). T cell cytolytic capacity is independent of initial stimulation strength. Nat Immunol 19, 849–858.

Rooney, M.S., Shukla, S.A., Wu, C.J., Getz, G., and Hacohen, N. (2015). Molecular and genetic properties of tumors associated with local immune cytolytic activity. Cell 160, 48–61.

Sharma, P., Hu-Lieskovan, S., Wargo, J.A., and Ribas, A. (2017). Primary, Adaptive, and Acquired Resistance to Cancer Immunotherapy. Cell 168, 707–723.

Shimizu, K., Sugiura, D., Okazaki, I.M., Maruhashi, T., Takegami, Y., Cheng, C., Ozaki, S., and Okazaki, T. (2020). PD-1 Imposes Qualitative Control of Cellular Transcriptomes in Response to T Cell Activation. Mol Cell 77, 937–950 e936.

Singer, M., Wang, C., Cong, L., Marjanovic, N.D., Kowalczyk, M.S., Zhang, H., Nyman, J., Sakuishi, K., Kurtulus, S., Gennert, D., et al. (2016). A Distinct Gene Module for Dysfunction Uncoupled from Activation in Tumor-Infiltrating T Cells. Cell 166, 1500–1511 e1509.

Stephens, M. (2017). False discovery rates: a new deal. Biostatistics 18, 275–294.

Subach, F.V., Subach, O.M., Gundorov, I.S., Morozova, K.S., Piatkevich, K.D., Cuervo, A.M., and Verkhusha, V.V. (2009). Monomeric fluorescent timers that change color from blue to red report on cellular trafficking. Nat Chem Biol 5, 118–126.

Trefzer, A., Kadam, P., Wang, S.H., Pennavaria, S., Lober, B., Akcabozan, B., Kranich, J., Brocker, T., Nakano, N., Irmler, M., et al. (2021). Dynamic adoption of anergy by antigen-exhausted CD4(+) T cells. Cell Rep 34, 108748.

Tubo, N.J., and Jenkins, M.K. (2014). TCR signal quantity and quality in CD4(+) T cell differentiation. Trends Immunol 35, 591–596.

Tumeh, P.C., Harview, C.L., Yearley, J.H., Shintaku, I.P., Taylor, E.J., Robert, L., Chmielowski, B., Spasic, M., Henry, G., Ciobanu, V., et al. (2014). PD-1 blockade induces responses by inhibiting adaptive immune resistance. Nature 515, 568–571.

Xiao, Z., Mayer, A.T., Nobashi, T.W., and Gambhir, S.S. (2020). ICOS Is an Indicator of T-cell-Mediated Response to Cancer Immunotherapy. Cancer Res 80, 3023–3032.

Yu, G., Wang, L.G., Han, Y., and He, Q.Y. (2012). clusterProfiler: an R package for comparing biological themes among gene clusters. OMICS 16, 284–287.

Yu, G., Wang, L.G., Yan, G.R., and He, Q.Y. (2015). DOSE: an R/Bioconductor package for disease ontology semantic and enrichment analysis. Bioinformatics 31, 608–609.

Zinzow-Kramer, W.M., Weiss, A., and Au-Yeung, B.B. (2019). Adaptation by naive CD4(+) T cells to self-antigen-dependent TCR signaling induces functional heterogeneity and tolerance. Proc Natl Acad Sci U S A 116, 15160–15169.

