## Supplementary Figures 1-5 for "Antigen and Checkpoint Receptor Recalibration of T Cell Receptor Signal Strength"

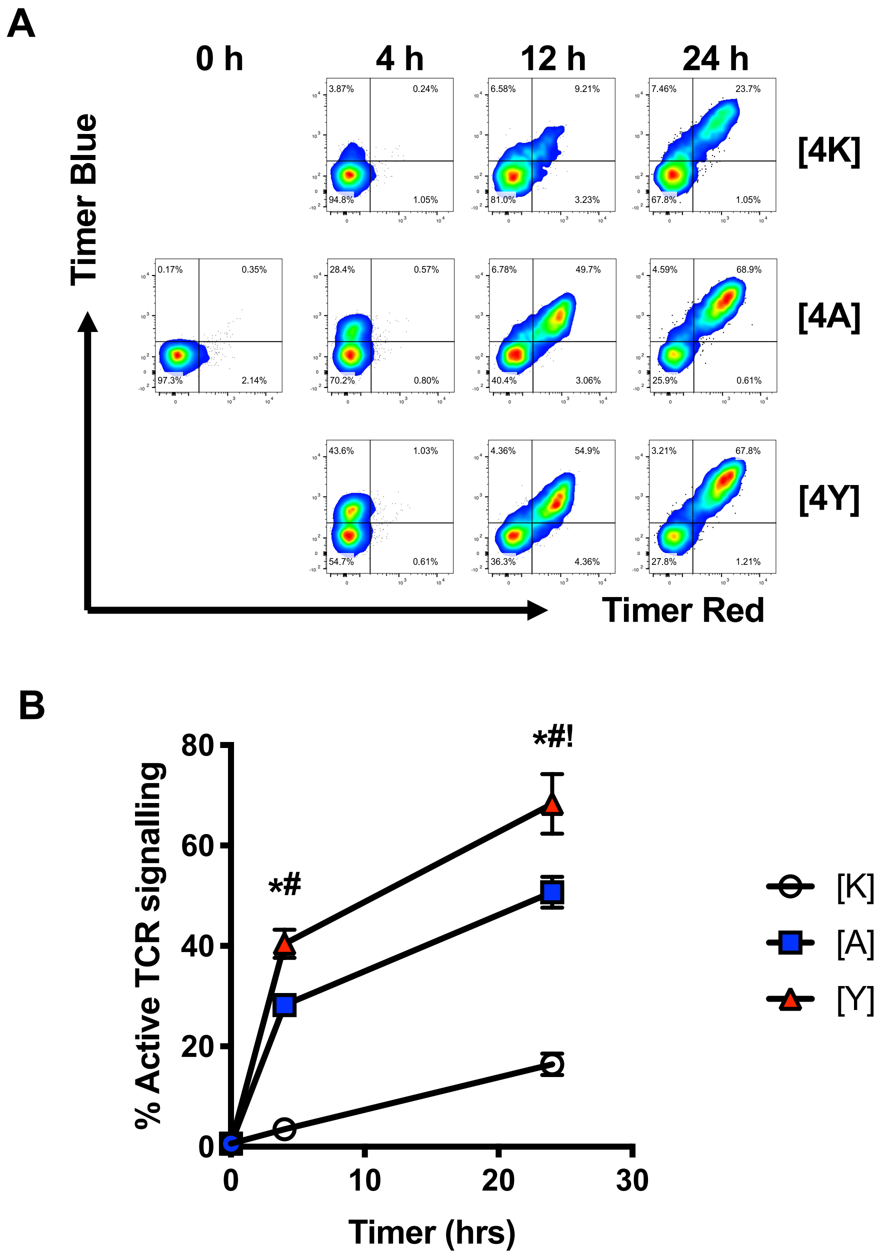


**Supplementary Figure 1: Modified MBP peptide variants induce potent T cell activation in vitro (related to Figure 1)**

**(A)** Naïve CD4+ T cells from Tg4 Nr4a3-Tocky IL10-GFP mice were incubated with CD90-depleted splenocytes in the presence of 1 μM of native [4K] MBP peptide, or [4A] or [4Y] variants for the times indicated before analysis of Nr4a3-Blue vs Nr4a3-Red expression in $CD4^{+}$ Tg4 T cells. (**B**) Summary data showing the % Nr4a3-$Blue^{+}$ in $CD4^{+}$ T cells in the 3 peptide groups. MBP [4K] = white circles, MBP [4A] = black squares and MBP [4Y] = red circles. Bars represent mean±SEM, n=3. Statistical analysis by two-way ANOVA with Tukey's multiple comparisons test. Significant differences between [4Y] and [4K] = *, [4A] and [4K] = #, or [4Y] and [4A] = !.

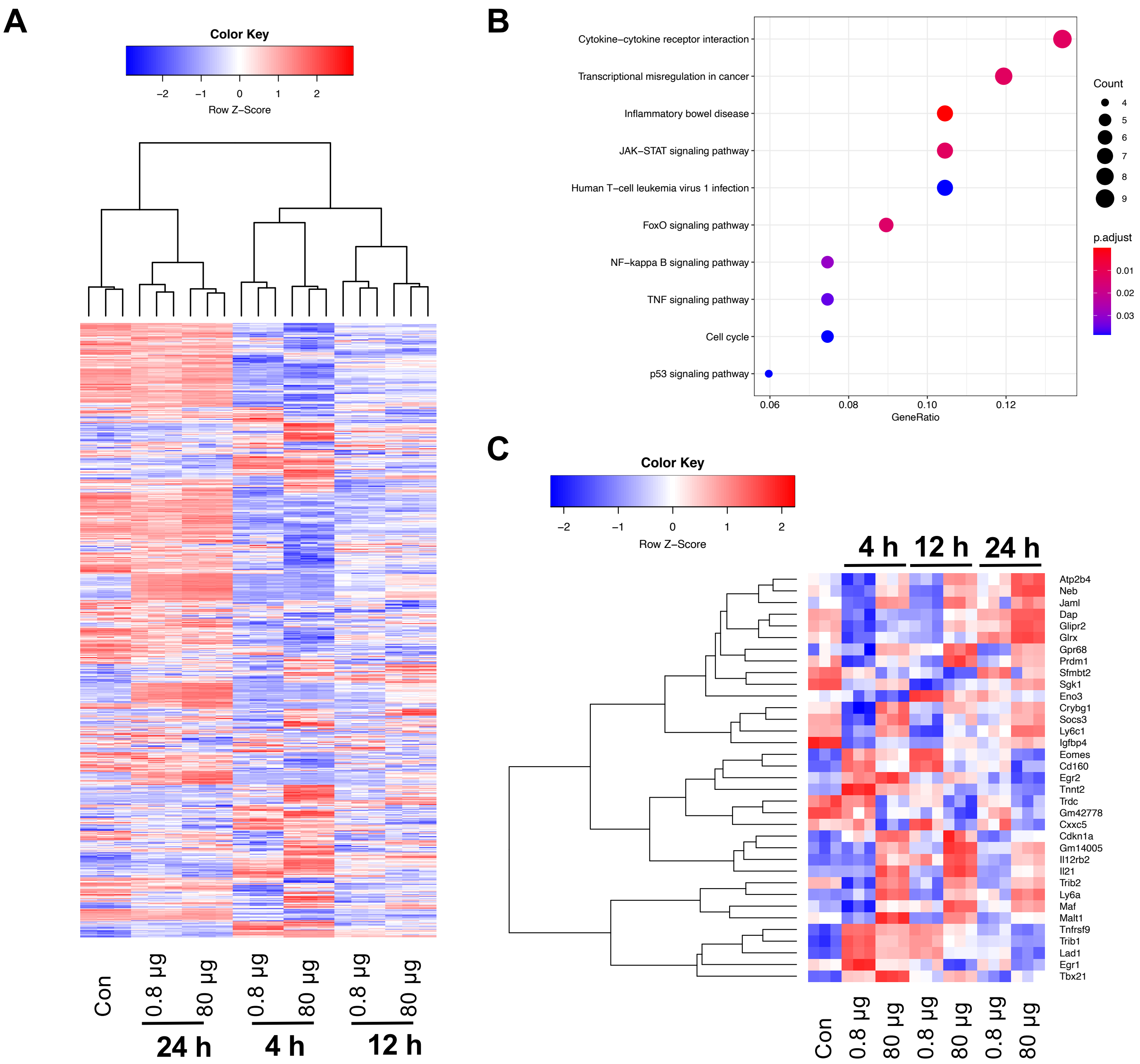


**Supplementary Figure 2: Analysis of DEGs in T cells receiving strong or weak TCR signalling in vivo (related to Figure 2).**

**(A)** Z-score heatmap analysis of log2 transformed and normalised counts for all unique DEG identified between 0.8 and 80 µg groups at 4, 12 or 24 h. (**B**) KEGG pathway analysis of DEG at 24 h between 0.8 µg and 80 µg [4Y] MBP immunised mice. (**C**) Z-score heatmap analysis of log2 transformed and normalised counts for genes that show differential expression across all 3 time points (4, 12 and 24 h) analysed between 0.8 µg and 80 µg immunised groups.

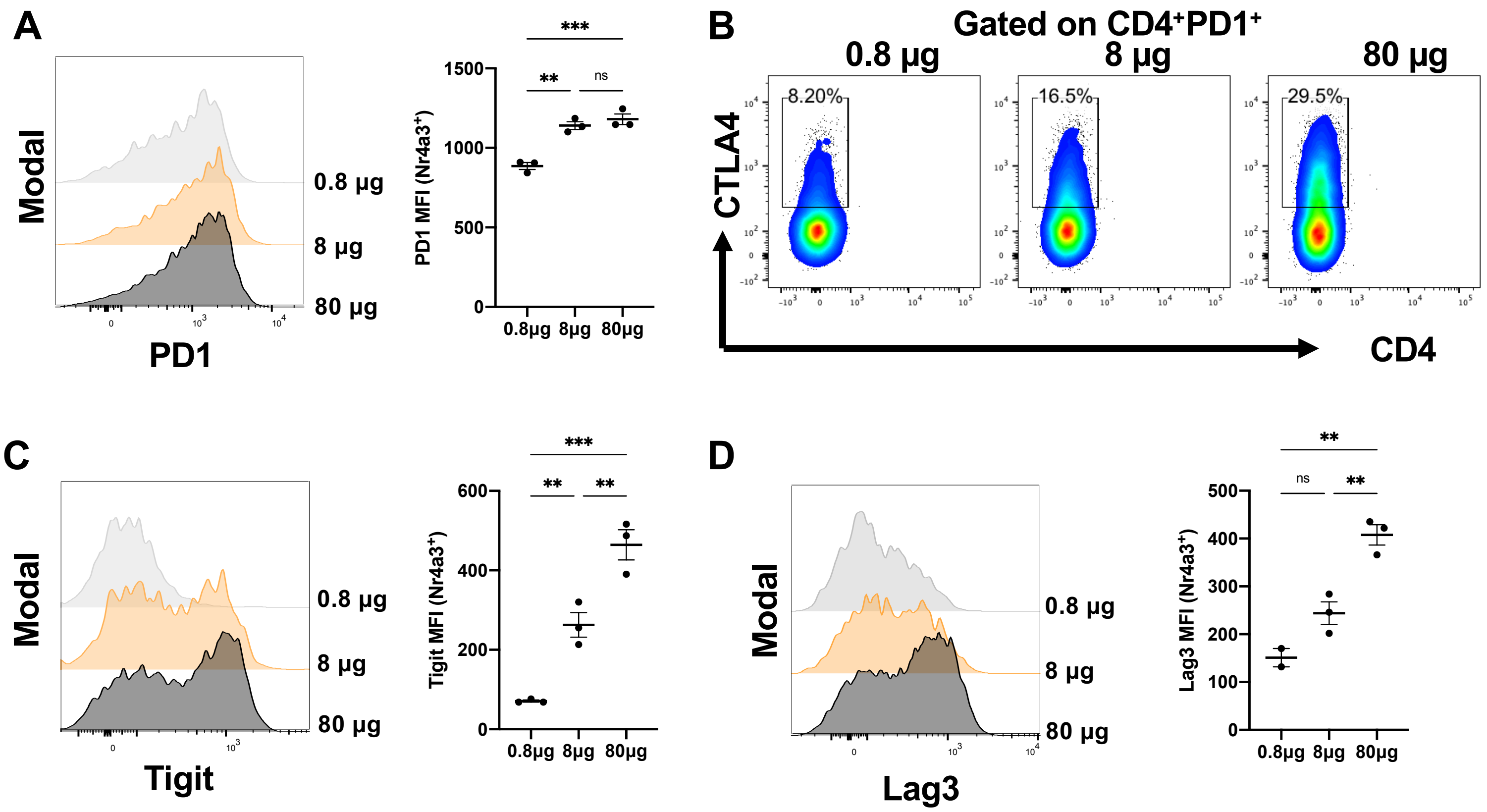


**Supplementary Figure 3: Protein expression patterns of PD1, CTLA4, Lag3 and TIGIT in response to different grades of TCR signal strength (related to Figure 3)**

**(A)** Tg4 Nr4a3-Tocky IL10-GFP mice were immunised s.c. with 0.8 µg, 8 µg or 80 µg of [4Y] MBP peptide and splenic CD4$^{+}$ Nr4a3-Timer$^{+}$ T cells (or gated on CD4$^{+}$ PD1$^{+}$ in (**B**)) analysed for PD1 (**A**), CTLA4 (**B**), Tigit (**C**) or Lag3 (**D**) expression. Dots represent individual mice and bars represent mean ±SEM. Statistical analysis by one-way ANOVA with Tukey's multiple comparisons test.

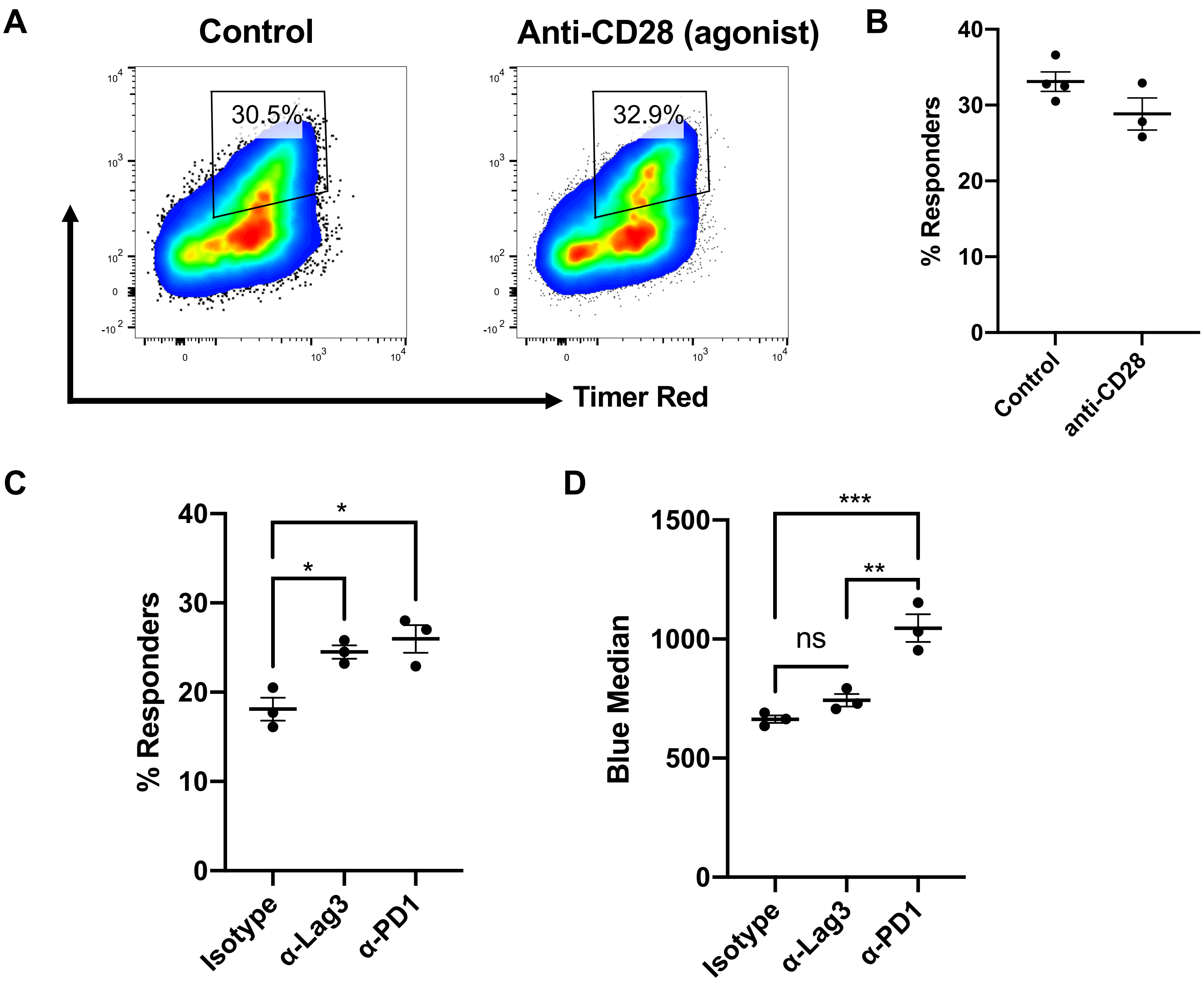


**Supplementary Figure 4: Effects of CD28, Lag3 and PD1 pathways on T cell re-activation in vivo (related to Figure 4)**

**(A)** Tg4 Nr4a3-Tocky IL10-GFP mice were immunised s.c. with 80 µg of [4Y] MBP. 24 h later mice were randomised to receive either PBS or agonistic anti-CD28 30 minutes prior to re-challenge with 8 µg [4Y] MBP peptide. Splenic $CD4^+$ T cells were analysed for Nr4a3-Timer Blue vs. Nr4a3-Timer Red analysis 4 h after peptide rechallenge. (**B**) Summary data of (**A**), control n=4, anti-CD28 n=3. (**C&D**) Tg4 Nr4a3-Tocky IL10GFP mice were immunised s.c. with 80 µg of [4Y] MBP. 24 h later mice were randomised to receive either isotype, anti-Lag3 or anti-PD1 30 minutes prior to re-challenge with 8 µg [4Y] MBP peptide. The frequency of responder (Nr4a3-$Blue^+Red^+$) T cells (**C**) or Nr4a3-Blue Median expression in responder (Nr4a3-$Blue^+Red^+$) T cells (**D**) 4 h after peptide rechallenge are shown. N=3, dots represent individual mice, bars represent mean ±SEM. Statistical analysis by one-way ANOVA with Tukey's multiple comparisons test. *=p<0.05, **=p<0.01, ***=p<0.001

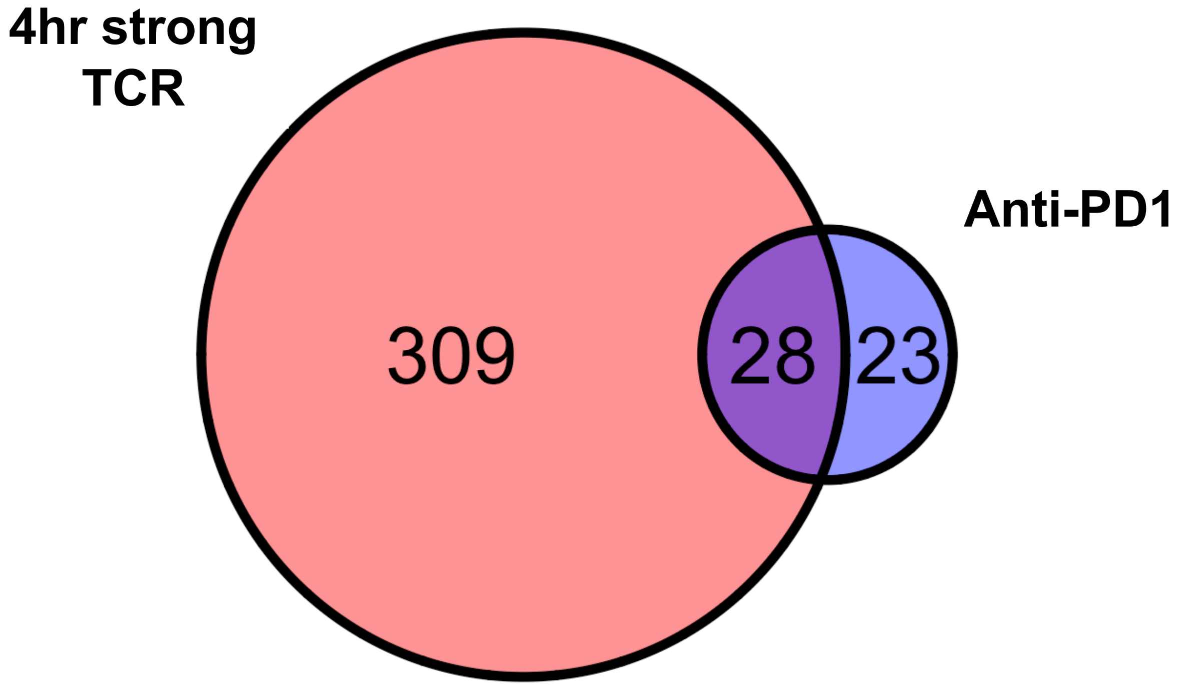


**Supplementary Figure 5: Majority of anti-PD1 specific T cell genes are upregulated in T cells receiving strong TCR signal (related to Figures 2,4 and 5)**

Genes upregulated in Tg4 CD4$^+$ T cells 4 h after receiving 80 µg vs 0.8 µg [4Y] MBP (Figure 2) were intersected with genes selectively upregulated at 4hrs of T cells re-activated in the presence of anti-PD1 in vivo (Figure 4). 28 out of 51 genes were overlapping.
